# Targeting of ibrutinib resistance–driving pathways by miR-28 in ABC-DLBCL

**DOI:** 10.1101/2025.11.12.687947

**Authors:** Emigdio Álvarez-Corrales, Rocío Moreno-Palomares, Carmen Gómez-Escolar, Mario Martínez, Udane Moral-Pérez, María Laguna-Herrero, Teresa Fuertes, Belén S. Estrada, Sonia Mur, Adriana de Bonis, Magdalena Leiva, Nuria Martínez-Martín, Álvaro Somoza, Almudena R. Ramiro, Virginia G. de Yébenes

## Abstract

Diffuse large B-cell lymphoma (DLBCL) is the most common aggressive B-cell lymphoma. Although many patients respond well to R-CHOP immunochemotherapy, those with the activated B-cell (ABC) subtype are often refractory or relapse. Bruton tyrosine kinase (BTK) inhibitors such as ibrutinib have improved outcomes, but acquired resistance limits their long-term efficacy. Here, we modeled the development of ibrutinib resistance in ABC-DLBCL and investigated whether the BCR-signaling regulator microRNA-28 (miR-28) can block this process. Using flow cytometry–based competition assays, multicolor clonal barcoding, transcriptomic profiling, and xenograft models, we found that miR-28 expression impairs the emergence of ibrutinib-resistant ABC-DLBCL cells. Mechanistically, miR-28 interferes with the clonal selection process triggered by ibrutinib treatment and rewires transcriptional programs by downregulating mitochondrial and mTOR signaling pathways critical for resistance development. Furthermore, the miR-28–repressed gene signature associated with ibrutinib resistance correlates with improved survival in ibrutinib-treated patients from the PHOENIX trial cohort with the MCD genetic subtype, which is associated with ABC-DLBCL. Finally, the targeted therapeutic delivery of miR-28 via aptamer-guided nanoparticles suppresses ibrutinib-resistant tumor growth *in vivo*. These findings identify miR-28 as an effective inhibitor of ibrutinib resistance, underscoring its translational potential as an adjunct strategy in ABC-DLBCL therapy.

**Graphical Abstract:** 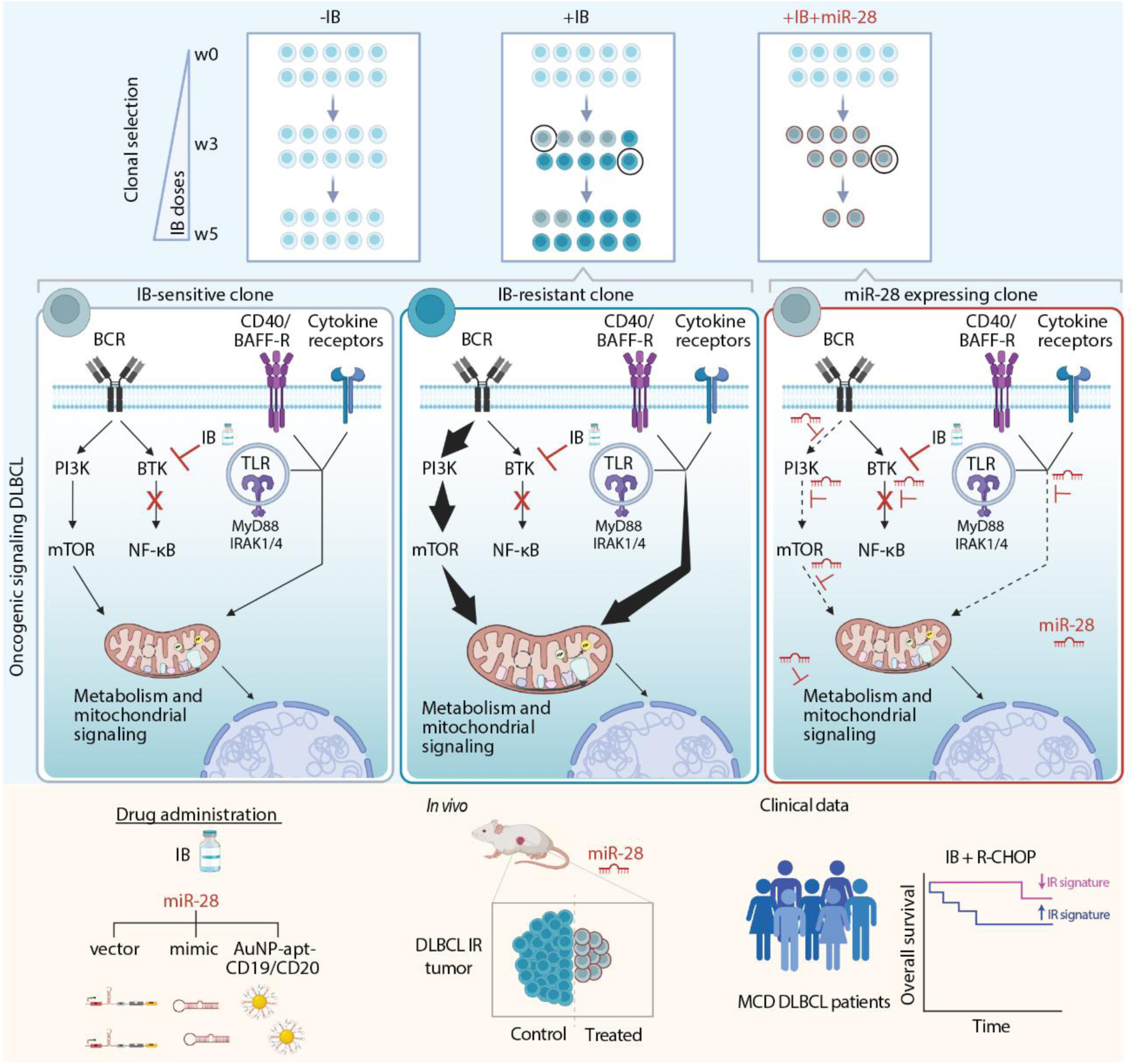

## INTRODUCTION

Diffuse large B-cell lymphoma (DLBCL) is the most common aggressive B-cell lymphoma(^1^). DLBCL patients are generally treated with a chemotherapy-based regimen consisting of rituximab plus cyclophosphamide, doxorubicin, vincristine, and prednisone (R-CHOP). Recent studies have aimed to define the molecular heterogeneity of DLBCL to support the development of targeted therapies. Based on gene expression profiling, DLBCL is classified into two main subtypes: germinal center B cell-like (GCB) and activated B cell-like (ABC), the latter is characterized by the expression of genes induced upon B-cell receptor (BCR) activation and is associated with a worse prognosis than GCB-DLBCL(^2^). Genetic profiling has identified distinct subtypes within ABC-DLBCL—primarily MCD, BN2, and N1—which differ not only in their genetic alterations but also in clinical outcomes(^3^). Overall survival after R-CHOP chemotherapy is achieved in only 40% of ABC-DLBCL patients and in as few as 26% of those with the MCD genetic subtype(^3, 4^). Functional genomic screens have identified genes essential for oncogenic signaling in DLBCL, including genes encoding proximal BCR signaling components as well as genes involved in distinct downstream survival pathways such as PI3K/AKT/mTOR and NF-κB. These screens have also revealed that ABC-DLBCL, but not GCB-DLBCL, is highly dependent on BCR-mediated NF-κB signaling for survival(^5–8^).

The introduction of Bruton’s tyrosine kinase (BTK) inhibitors, such as ibrutinib, which blocks BCR-dependent NF-κB activation, has been transformative in the treatment of several B-cell malignancies(^9^). Clinical trials have also demonstrated the potency of BTK inhibition, either as monotherapy or in combination with R-CHOP, in ABC-DLBCL(^10–14^). However, responses to ibrutinib in ABC-DLBCL are variable or incomplete across different age groups and genetic subtypes, and acquired resistance remains a significant barrier to achieving durable outcomes(^10, 14^). Prolonged ibrutinib treatment has been associated with the emergence of resistance, and mutations affecting BTK, its downstream effector PLCG2, or CARD11 have been identified in CLL and DLBCL patients(^10, 15–18^). Importantly, beyond these genetic alterations, resistance also emerges through non-genetic mechanisms, such as rewiring of oncogenic signaling pathways and feedback activation of compensatory survival programs(^19–21^). Addressing these adaptive responses to ibrutinib treatment requires rational combinations of targeted therapies designed to inhibit different resistance-promoting pathways simultaneously.

In this context, microRNAs (miRNAs) have gained increasing attention as potential therapeutics for B-cell malignancies. miRNAs are small non-coding RNAs that regulate gene expression post-transcriptionally by binding to target messenger RNAs (mRNAs), resulting in mRNA degradation or translational inhibition. Each miRNA can modulate the expression of multiple mRNAs, thereby influencing complex gene regulatory networks(^22^). During oncogenesis, miRNA-mediated regulation is frequently disrupted. Several miRNAs have been identified as key contributors to B-cell lymphoma pathogenesis, functioning as either tumor suppressors or oncogenic miRNAs(^23^).

Recent advances have enabled the modulation of miRNA activity through the delivery of synthetic mimics or inhibitors, offering new therapeutic opportunities for various diseases, including cancer. In this context, miRNA-based therapies have emerged as promising alternatives to conventional treatments for B-cell non-Hodgkin lymphoma (B-NHL) and other malignancies(^24–26^). A key advantage of these strategies is their ability to simultaneously target multiple components of oncogenic signaling pathways, potentially limiting the emergence of resistance mechanisms such as mutations in individual oncogenes.

miR-28 is a microRNA with a downregulated expression in B-cell lymphomas(^27^). We previously reported that it targets a BCR-dependent signaling network in B cells and exerts anti-tumor effects in DLBCL, both as a single agent(^28^) and in combination with ibrutinib, by promoting cell-cycle arrest and impairing DNA replication(^29^). In the present study, we evaluated the role of miR-28 in preventing the emergence of ibrutinib resistance. Our findings show that miR-28 disrupts the post-transcriptional adaptation and clonal selection processes that occur during the development of resistance to ibrutinib in ABC-DLBCL. Moreover, we describe the synthesis and *in vivo* antitumor efficacy of CD19/CD20-aptamer–functionalized gold nanoparticles (AuNPs) that efficiently deliver a miR-28 mimic to resistant DLBCL cells, achieving sustained tumor growth suppression.

## MATERIALS AND METHODS

### Lentiviral expression constructs

Two types of lentiviral vectors were used: pTRIPZ: doxycycline-inducible constructs for the expression of either miR-28 or a scramble control sequence and LeGO vectors (LeGO-V2, LeGO-Cer2): applied for multicolor genetic labeling in cell tracking assays.

### RNA sequencing

Bulk RNA sequencing was performed on the following clones from MD-901 color-barcoded cells (CBC) after three weeks of exposure to gradually increasing doses of ibrutinib (ranging from 10-fold below to 2.5-fold above the GR_50_): ibrutinib-sensitive (scr week 5 < week 1, n = 4), ibrutinib-resistant (scr week 5 > week 1, n = 4), and miR-28–expressing (n = 5).

### Lymphoma models

For subcutaneous xenografts, a suspension of 1–5 × 10⁶ ABC-DLBCL cells was prepared in 100 μL of phosphate-buffered saline (PBS), mixed 1:1 with Matrigel (BD Biosciences), and injected into the flanks of 10–12-week-old NSG mice.

### Treatment with aptamer-conjugated AuNP-miR-28

miR-28- or scr-functionalized nanoparticles conjugated with CD19 or CD20 aptamers were used to treat established IR ABC-DLBCL tumors (>150 mm³). Tumors were treated twice weekly with intratumoral (2.3 pmol) or intravenous (23 pmol) injections of nanoparticles, containing either miR-28 or the control sequence.

Further methodological details are provided in the Supplementary Material.

## RESULTS

### miR-28 inhibits the generation of ibrutinib resistance in ABC-DLBCL cells

To evaluate the impact of miR-28 on the development of ibrutinib resistance in ABC-DLBCL, we first established a polyclonal competition assay using flow cytometry and assessed the effect of miR-28 in four human ABC-DLBCL cell lines: MD-901, U2932, Riva, and HBL1. In this assay, we mixed, in a 1:1 ratio, populations of each ABC-DLBCL cell line transduced with pTRIPZ doxycycline-inducible lentiviral vectors encoding either miR-28 or a scramble (scr) sequence, along with RFP(^28^). The pTRIPZ-transduced ABC-DLBCL cells used in the assay were additionally labeled by transduction with vectors encoding fluorescent proteins, Cerulean for miR-28-transduced cells and Venus for scr-transduced cells (Figure 1A). The mixed miR-28-Cerulean and scr-Venus populations were cultured for five weeks in the presence of doxycycline and exposed to ibrutinib at concentrations that increased weekly from approximately 10-fold below to approximately 2-fold above the ibrutinib growth rate inhibition 50% value (GR_50_). This stepwise drug escalation protocol results in the generation of ibrutinib-resistant ABC-DLBCL cells (Figure S1A–B), as previously described(^20^), which can be stably maintained *in vitro* at 2×GR_50_ ibrutinib. Analysis of miR-28 expression in the four non-transduced ABC-DLBCL cell lines, which exhibit markedly reduced miR-28 levels compared with non-tumoral B cells(^29^), showed no significant changes after the resistance-generation process (Figure S1C). The mixed miR-28-Cerulean and scr-Venus cultures were analyzed weekly by flow cytometry to quantify changes in the relative proportions of miR-28-Cerulean and scr-Venus cells (Figure 1A-B). We found that the proportion of RFP^+^ miR-28-Cerulean^+^ cells in the four ABC-DLBCL cell lines gradually decreased as ibrutinib concentrations increased, and that the contribution of miR-28-expressing cells in ibrutinib-resistant cultures (week 5) was significantly lower than in the starting cultures (Figure 1B-C). RFP^+^ miR-28-Cerulean^+^ cells accounted for a minor proportion of the cells (18% of U2932, and ≤ 9% of MD-901, Riva and HBL1) after five weeks of culture in the presence of ibrutinib (Figure S1D). To exclude any possible effect of Cerulean and Venus fluorescent protein expression on the competition assay, we also performed a control competition assay in scr-transduced MD-901 ABC-DLBCL cells. As expected, we found no significant differences in the proportions of scr-Cerulean and scr-Venus populations after five weeks of culture with increasing ibrutinib concentrations (Figure 1D, S1E). In addition, we analyzed how miR-28 expression affects cell fitness in competition assays in the absence of ibrutinib. Five-week cell-competition assays performed in all four ABC-DLBCL cell lines cultured without ibrutinib showed a variable decrease in the proportions of miR-28–expressing cells, which was greater in HBL1 and U2932 than in MD-901 and Riva cells (Figure S2A). Notably, the loss of miR-28–expressing cells was more pronounced across all cell lines in the presence of increasing ibrutinib concentrations, indicating that the strong negative selection against miR-28–expressing cells is largely driven by ibrutinib exposure (Figure S2A). In agreement with the previous results, we found that the generation of ibrutinib-resistant cells was diminished in MD-901, U2932, and HBL1 ABC DLBCL cells expressing miR-28 (Figure S2B), and that miR-28 pTRIPZ-transduced Riva ABC DLBCL cell cultures were enriched in RFP^-^ cells by the time when ibrutinib-resistant generation had occurred (week 5) (Figure S2C). Together, these results show that miR-28 inhibits the generation of ibrutinib-resistant ABC-DLBCL cells.

**Figure 1.**
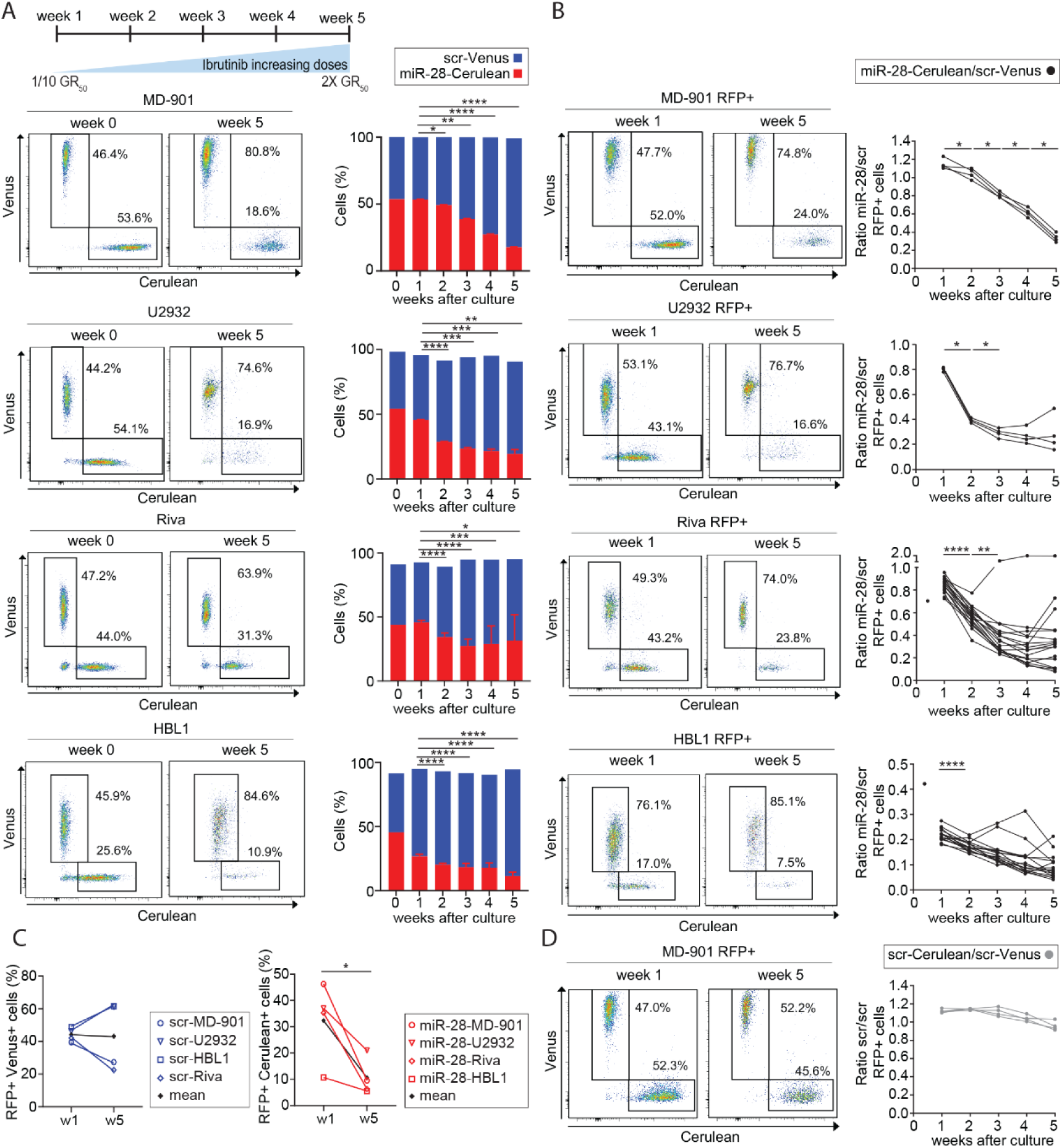
miR-28 limits the generation of ibrutinib resistance in polyclonal DLBCL cell competition assays. Polyclonal ibrutinib-resistance competition assays were performed by mixing ABC-DLBCL cells transduced with doxycycline-inducible RFP expressing lentiviral vectors encoding either miR-28 or a scramble (scr) sequence and labeled with either Cerulean or Venus fluorescent proteins at a 1:1 ratio. Cultures were exposed to ibrutinib for five weeks, with the drug concentration increasing weekly to reach up to 2-fold above the ibrutinib GR_50._ **(A–B)** Representative flow cytometry plots of scr-Venus/miR-28-Cerulean co-cultures and their evolution over five weeks in MD-901, U2932, Riva, and HBL1 cell lines. The bar chart (A) shows the progression of the total miR-28-Cerulean and scr-Venus populations over time (fold change of miR-28-Cerulean cells (week 5 versus week 1): 0.33 MD-901, 0.43 U2932, 0.69 Riva and 0.42 HBL1). The graphs on the right (B) display the growth ratio of miR-28-Cerulean to scr-Venus within the RFP⁺ population. **(C)** Percentage of RFP⁺ cells in co-cultures with either scr-Venus (left) or miR-28-Cerulean (right) at weeks 1 and 5 of cell competition in the presence of ibrutinib, across four DLBCL cell lines. (**D)** Cell competition assay of scr-Cerulean/scr-Venus MD-901 cells cultured with increasing doses of ibrutinib for five weeks. Representative flow cytometry plots of the co-cultures (left) and percentage of MD-901 RFP⁺ populations over time (right). Statistical analysis: One-way ANOVA analysis of changes in miR-28–Cerulean populations, with comparisons at each time point performed relative to week 1 (A), unpair *t*-test (B-D). \**P*<0.05, \*\**P*<0.01, ***P<0.001,\*\*\*\**P*<0.0001.

### miR-28 inhibits clonal selection during ibrutinib resistance generation in ABC-DLBCL cells

To determine whether miR-28 limits the generation of ibrutinib-resistant cells solely by impairing proliferation or promoting cell death, or whether it can actively suppress the development of resistance, we developed a flow cytometry–based assay to analyze resistance emergence at the clonal level using MD-901 cells. We first assessed whether ABC-DLBCL ibrutinib resistance competition assays could be developed with clonal resolution by plating 10 miR-28–Cerulean and scr–Venus MD-901 cells mixed at a 1:1 ratio and culturing them for five weeks under escalating ibrutinib exposure. Our results confirmed the loss of miR-28–expressing cells in this setting as well (Figure S3A). Next, to monitor clonal diversity, we generated a color-barcoded (CBC) ABC-DLBCL cell population by transducing MD-901 cells with lentiviral vectors encoding Venus and Cerulean fluorescent proteins. Ten CBC cells, each expressing a random number of Venus and/or Cerulean-encoding lentiviral copies, were seeded into 96-well plates. These fluorescently barcoded cells were then treated for five weeks with ibrutinib at concentrations that increased weekly from 10-fold below to 2-fold above the GR_50_ in MD-901 cells, or were left untreated (n = 45 wells per condition). The proportions of the different CBC clones in each well were monitored weekly by flow cytometry to quantify changes over time. We found that the gradual increase in ibrutinib concentration led to the selection of a subset of clones—approximately 25%—within each well (Figure S3B), whereas the proportions of clones in untreated wells remained more stable over time (Figure 2A-B). In ibrutinib-treated wells, cells derived from ibrutinib-selected clones expanded to constitute roughly 75% of the culture (Figure S3C), resulting in a gradual reduction of diversity index of ibrutinib-treated wells over time compared to untreated wells (Figure 2C; S3D). Thus, we conclude that the CBC assay enables monitoring of clonal selection and the emergence of ibrutinib-resistant clones that outcompete ibrutinib-sensitive cells during the development of resistance in ABC-DLBCL cells.

**Figure 2.**
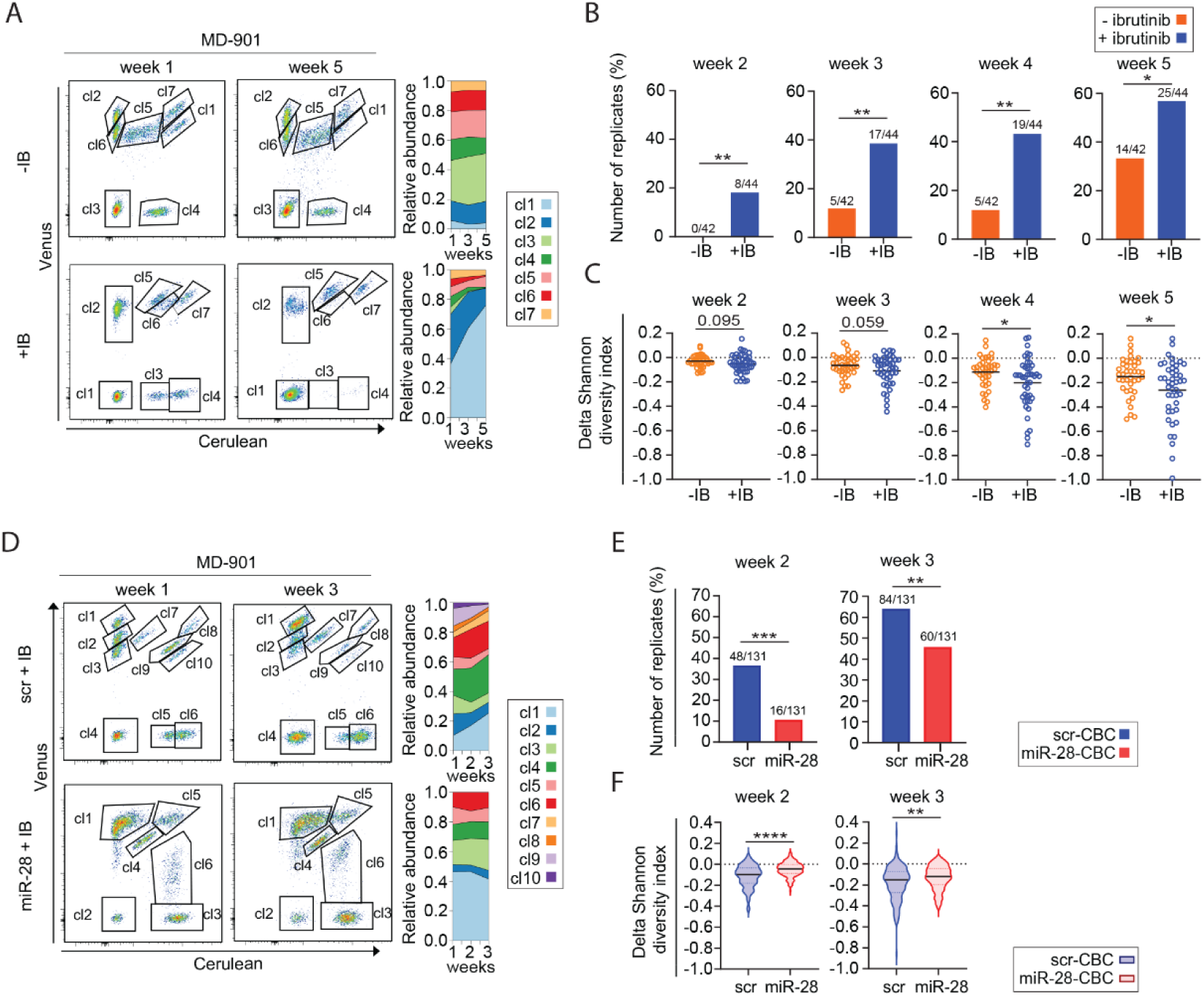
miR-28 inhibits clonal selection during acquisition of ibrutinib resistance in DLBCL cells. Color-barcoded MD-901 ABC-DLBCL cells (CBC), each carrying a random number of Venus- and/or Cerulean-encoding lentiviral constructs, were seeded into 96-well plates at an average density of 10 cells per well and cultured for up to five weeks. In panels A–C, cells were either exposed to gradually increasing concentrations of ibrutinib (IB), with doses escalating weekly from 10-fold below to 2-fold above the GR_50_, or left untreated (±IB). In panels D–F, CBC cells transduced with doxycycline-inducible RFP-expressing lentiviral vectors encoding either miR-28 or a scramble control (scr) were cultured under increasing ibrutinib concentrations (+IB) to assess the impact of miR-28 expression on clonal dynamics during acquisition of drug resistance (n = 45 wells per condition). **(A)** Representative wells showing clonal composition after one (left) or five (right) weeks under ibrutinib (+IB; well 1C) or control (-IB; well 11C) conditions, analyzed by flow cytometry. Clone proportions in these wells over time are shown in the Muller plot on the right. **(B)** Percentage of wells with changes in clone proportions compared to week one under +IB (blue) or -IB (orange) conditions. A significant change was defined as a >3-fold decrease at weeks 2–3, and >5-fold at weeks 4–5. **(C)** Clonal diversity measured by the Shannon index in wells treated with ibrutinib (blue) or left untreated (orange). Relative diversity per week, normalized to week 1 (C). Error bars represent standard deviation (n = 45 wells/condition). **(D)** Representative wells showing clonal composition after one (left) or three (right) weeks in scramble (scr+IB; well 9A) or miR-28 (miR-28+IB; well 5D) cultures, analyzed by flow cytometry. Clone proportions over time in each well are shown in the Muller plots on the right. **(E)** Percentage of wells with significant changes in clone proportions compared to week 1 in scr (blue) or miR-28 (red) conditions. A significant change was defined as a >3-fold decrease. **(F)** Clonal diversity over time measured by the Shannon index, normalized to week 1, in scr (blue) and miR-28 (red) CBC wells. Data pooled from three independent experiments (n=45 wells/experiment and condition; E-F). Statistical analysis: Fisher’s exact test (B and E), unpaired t-test (C and F). \**P* < 0.05; \*\**P* < 0.01; \*\*\**P* < 0.001; \*\*\*\**P* < 0.0001.

Next, we used the CBC clonal assay to test the effect of miR-28 expression on the generation of ibrutinib resistance. We found that miR-28 expression impaired the emergence of ibrutinib-resistant clones, as the proportions of CBC clones remained more stable during the first three weeks of culture, with gradually increasing ibrutinib doses, in wells seeded with miR-28-expressing CBC cells compared to those seeded with cells expressing a scramble construct (Figure 2D-E). Accordingly, the diversity index of miR-28 CBC wells was higher than that of scramble CBC wells (Figure 2F).

After five weeks of culture, when ibrutinib concentration is 2-fold above the MD-901 GR_50_, we found that scramble CBC wells reached a high cellular density, whereas wells containing miR-28 CBC cells were more heterogeneous in cell density and included medium and low cellular densities (Figure S3E). RFP expression analysis revealed that wells with low cellular density had a higher percentage of RFP^+^ cells than medium-density miR-28 CBC wells (Figure 3A). Moreover, t-SNE analysis of RFP expression showed that miR-28 RFP^+^ CBC cells present at week 1 were preferentially depleted at week 5 in medium-density wells, but not in low-density wells (Figure 3B). Sorting miR-28-pTRIPZ–transduced MD-901 cells by RFP levels confirmed that RFP intensity reflects miR-28 expression levels (Figure S3F). These results suggest an inverse correlation between RFP expression and cell survival of miR-28–expressing cells in gradually increasing ibrutinib cultures. To determine whether there is a correlation between RFP expression and the generation of ibrutinib-resistant cells in miR-28 CBC wells, we quantified the RFP mean fluorescent intensity of all the CBC clones (655 miR-28 CBC clones and 617 scr CBC clones), analyzing them according to their relative increase or decrease in proportions in week 5 (w5) *versus* week 1 (w1) cultures. We found that RFP expression was higher in miR-28 CBC clones that were counter-selected in the presence of ibrutinib, but did not change in scramble CBC clones regardless of whether the clones had become resistant to ibrutinib (w5 > w1) or were ibrutinib-sensitive (w5 < w1) (Figure 3C). In addition, we quantified the number of ibrutinib-resistant cells generated in scramble and miR-28 CBC wells. We found a reduced number of viable CBC cells after five weeks of culture with increasing ibrutinib concentrations in miR-28 wells (Figure 3D, S3G). Together, these results show that miR-28 impairs the generation of ibrutinib-resistant ABC-DLBCL cells.

**Figure 3.**
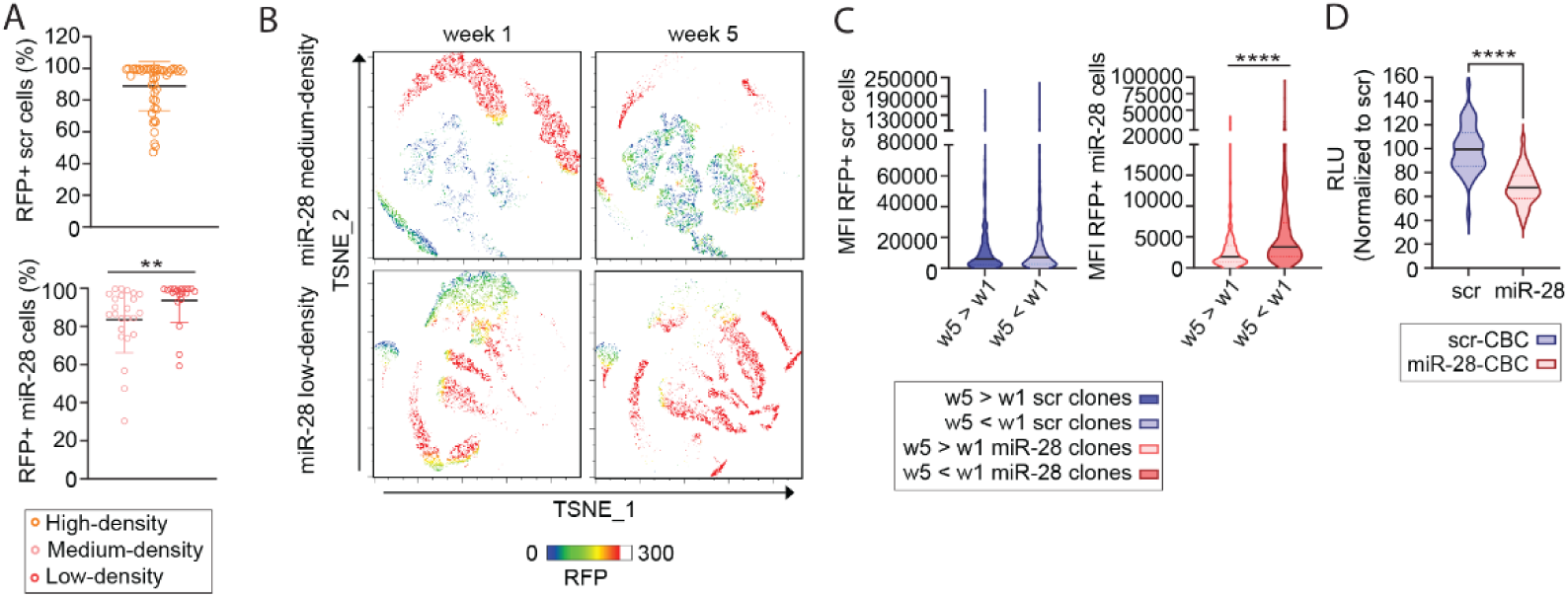
miR-28 interferes with the development of ibrutinib resistance in DLBCL cells. Color-barcoded MD-901 ABC-DLBCL cells (CBC) transduced with doxycycline-inducible, RFP-expressing lentiviral vectors encoding either miR-28 or a scramble control (scr) were seeded at an average of 10 cells per well and exposed to increasing concentrations of ibrutinib (IB), reaching 2-fold above the GR_50_ after five weeks (n = 45 wells/condition and experiment). **(A)** Quantification of the percentage of RFP⁺ cells in wells grouped by cellular density (high, orange; medium, pink; or low, red), as determined by culture color. Cell culture images and classification criteria are shown in Figure S3E. **(B)** Representative examples of medium-density (well 5C) and low-density (well 5D) wells from the miR-28 group. FACS data from weeks 1 and 5 were concatenated and analyzed by t-SNE. Color gradient indicates RFP intensity. **(C)** Quantification of RFP mean fluorescence intensity (MFI) per clone, categorized by increased (w5 > w1) or decreased (w5 < w1) abundance at week 5 relative to week 1. Data pooled from three independent experiments. **(D)** Viable cell numbers in scr and miR-28-expressing wells at week 5, determined using CellTiter-Glo®. Luminescence was expressed as Relative Luminescence Units (RLU) and normalized to the mean of scr wells. Error bars represent standard deviation. Statistical analysis: unpaired t-test (A, C and D). \*\**P* < 0.01 \*\*\*\**P* < 0.0001.

### miR-28 inhibits signaling pathways required for the development of ibrutinib resistance in ABC-DLBCL

To identify miR-28–regulated signaling pathways involved in the acquisition of ibrutinib resistance, we performed RNA transcriptomic analysis after three weeks of culture with gradually increasing doses of ibrutinib, prior to the full establishment of resistance. We analyzed scramble ibrutinib-sensitive clones (defined as week 5 < week 1), scramble ibrutinib-resistant clones (defined as week 5 > week 1), and miR-28–expressing CBC clones (Figure S4). Principal component analysis (PCA) of transcriptomic profiles revealed clustering by condition, with high intra-group similarity across the three experimental groups (Figure 4A). RNA-seq analysis comparing ibrutinib-sensitive (n = 4) and week 3–resistant scramble (n = 4) CBC clones identified 695 differentially expressed genes (DEG) associated with resistance development. Transcriptomic comparison between miR-28–expressing clones (n = 5) and week 3–resistant (n = 4) CBC clones revealed miR-28–dependent modulation of 3,370 transcripts (Table S1). In addition, comparison between miR-28–expressing clones (n = 5) and week 3–sensitive scramble clones (n = 4) identified 3,694 DEGs (Figure S5A and Table S1). To assess whether miR-28–mediated transcriptional changes reflect direct target regulation, we analyzed the distribution of evolutionarily conserved miR-28 binding sites among genes differentially expressed in week 3 CBC clones. Genes found downregulated in miR-28–expressing clones versus ibrutinib-resistant clones were significantly enriched for conserved miR-28 targets, whereas no enrichment was observed among upregulated genes (Figure S5B). Similarly, we observed a significant enrichment of miR-28 predicted targets among downregulated genes in miR-28–expressing MD-901 tumors from ibrutinib-treated mice (Figure S5C). Together, these results suggest that miR-28 regulates ABC-DLBCL gene expression through both direct and indirect effects on gene expression. Notably, Gene Set Enrichment Analysis (GSEA) showed that week 3–resistant CBC clones upregulated the expression of genes previously found to be essential for BTK-inhibitor resistance in HBL1 and TMD8 ABC-DLBCL cells using a genome-wide CRISPR-Cas9 screen(^20^) (p = 0.009, ES = 0.43) (Figure S5D). In contrast, the expression of these resistance-associated genes was significantly downregulated in miR-28–expressing CBC clones (p ≤ 0.000, ES = –0.46) (Figure S5D). Ingenuity Pathway Analysis (IPA) further revealed that resistance acquisition in scramble CBC clones is linked to downregulation of gene networks involved in cellular stress responses (e.g., eIF2 signaling, nonsense-mediated decay, amino acid deficiency response, ribosomal quality control) and different metabolic processes (e.g., translation, selenoamino acid metabolism, oxidative phosphorylation). Conversely, upregulation of mTOR and mitochondrial signaling was observed in week 3–ibrutinib resistant CBC clones (Figure S5E, Table S2). In miR-28–expressing CBC clones, these pathway alterations were reversed, with activation of stress and metabolic programs and suppression of mTOR and mitochondrial signaling (Figure S5E, Table S2). Consistently, GSEA using MitoCarta 3.0(^30^) confirmed that mitochondrial pathway expression was downregulated in miR-28–expressing CBC clones and upregulated in ibrutinib-resistant CBC clones following three weeks of ibrutinib exposure (Figure 4B). In agreement with these results, we found that mTOR and mitochondrial signaling pathways were inhibited in miR-28–expressing MD-901 tumors from ibrutinib-treated mice (Figure S5E-F). Importantly, analysis of publicly available datasets from ibrutinib-resistant HBL1, OCI-Ly10, and OCI-Ly1 models(^31^) also revealed upregulation of mitochondrial-related pathways linked to ibrutinib resistance (Figure S5F).

**Figure 4.**
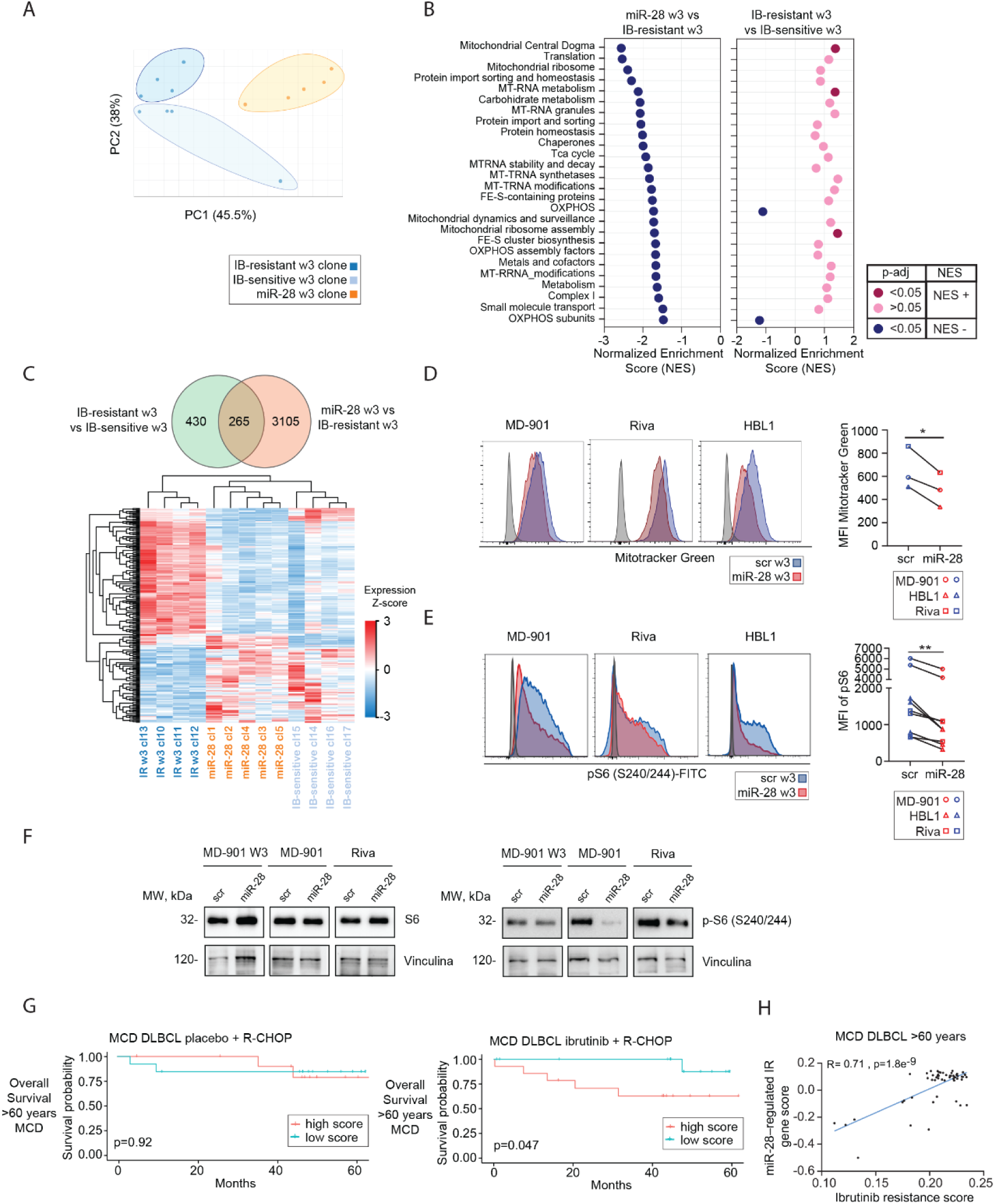
miR-28 suppresses pathways involved in ibrutinib resistance in ABC-DLBCL. **(A–C)** Transcriptomic analysis of sorted color-labeled MD-901 (CBC) clones after three weeks (w3) of culture with gradually increasing concentrations of ibrutinib (IB). CBC cells transduced with doxycycline-inducible, RFP-expressing lentiviral vectors encoding either miR-28 or a scramble control (scr) were seeded at an average density of 10 cells per well and exposed to IB, as described in Figure 2. **(A)** Principal component analysis of scr ibrutinib-resistant (IR) w3 clones (dark blue, n = 4), scr ibrutinib-sensitive w3 clones (light blue, n = 4), and miR-28–expressing (orange, n = 5) CBC clones. **(B)** GSEA of mitochondrial-related pathways (Mitocarta 3.0) in miR-28 and IR w3 CBC clones. Normalized enrichment scores (NES) and adjusted p-values (Benjamini–Hochberg correction) of altered pathways are shown in a scatter plot (blue dots, inhibited; pink dots, activated). **(C)** Heatmap showing hierarchical clustering based on Z-score expression values of the 265 genes involved in the acquisition of ibrutinib resistance and regulated by miR-28 in MD-901 clones. **(D–E)** Flow cytometry analysis of mitochondrial mass (D), assessed using MitoTracker Green staining, and S6 phosphorylation (Ser 240/244; E) in bulk miR-28- or scr-transduced MD-901, Riva, and HBL1 ABC-DLBCL cell lines after 3 weeks of culture in the presence of increasing doses of ibrutinib. Left: representative flow cytometry plots for each cell line (blue: scramble control; red: miR-28 overexpression; grey: unstained control). Right: Mean Fluorescence Intensity (MFI). Unpaired t-test; \**P* < 0.05; \*\**P* < 0.01. **(F)** Western blot analysis of S6 and p-S6 (Ser240/244). Cell lysates from bulk miR-28– or scramble-transduced MD-901 cells cultured for 3 weeks in the presence of doxycycline and increasing doses of ibrutinib (W3), as well as from sorted RFP^+^ MD-901 and Riva ABC-DLBCL cells two days after doxycycline administration, were analyzed. **(G)** Kaplan–Meier analysis of overall survival in MCD DLBCL patients aged ≥60 (n = 54), stratified by a scored miR-28–IR gene signature in baseline biopsies from the PHOENIX trial(^12^), comparing patients treated with R-CHOP and R-CHOP plus ibrutinib: highest 50^th^ percentile (up, red), and lowest 50^th^ percentile (down, green). \**P* < 0.05 (Log-rank Mantel–Cox test). **(H)** Correlation analysis of the miR-28–IR gene signature score and ibrutinib-resistance score in tumors from MCD DLBCL patients aged ≥60 years (n = 54) from the PHOENIX trial(^12^).

Next, we analyzed the specific set of genes involved in the acquisition of ibrutinib resistance that are regulated by miR-28 in CBC MD-901 clones, by comparing DEGs between scramble ibrutinib-resistant and ibrutinib-sensitive clones, as well as between miR-28–expressing and scramble ibrutinib-resistant clones at week 3. We found that miR-28 regulates 265 of the 695 DEGs associated with resistance development (Figure 4C). Hierarchical clustering of CBC clones based on the expression of these 265 genes showed that miR-28–expressing CBC clones clustered with ibrutinib-sensitive clones, but not with week 3–resistant clones. This clustering pattern reflects the coordinated regulation of this gene subset shared between miR-28–expressing and ibrutinib-sensitive clones, which is associated with the suppression of mTOR (Figure 4C, S5E, Table S2). We refer to this subset as the miR-28–regulated ibrutinib-resistance (miR-28-IR) gene signature. Given these transcriptomic findings, which reveal changes in cellular metabolism pathways related to mitochondrial biogenesis, we next assessed the mitochondrial status of scramble and miR-28–expressing ABC-DLBCL cells after three weeks of culture in the presence of gradually increasing doses of ibrutinib. MitoTracker Green staining and flow cytometry revealed reduced mitochondrial mass in miR-28–expressing MD-901, Riva, and HBL1 cells compared to scramble controls (Figure 4D). Seahorse analysis further showed that, although oxygen consumption rate (OCR) and extracellular acidification rate (ECAR) were similar between groups, miR-28–expressing MD-901 cells exhibited lower mitochondria-derived ATP production (Figure S6A). To evaluate mTOR activity, we measured the primarily mTORC1-dependent phosphorylation of ribosomal protein S6 (pS6) at Ser240/244 by flow cytometry. pS6 (Ser240/244) levels were consistently lower in miR-28–expressing MD-901, Riva, and HBL1 cells after three weeks of ibrutinib exposure (Figure 4E). These results were confirmed by immunoblotting in MD-901 cells after three weeks of ibrutinib treatment and in Riva and MD-901 cells after transient (two days) miR-28 expression (Figure 4F). Western blot analysis of predicted miR-28 targets upstream of mTOR signaling, including Akt2 (PI3K/Akt pathway) and Erk2 (Raf/ERK pathway), showed a mild miR-28–dependent downregulation of Akt2 in both MD-901 and Riva cells, and of Erk2 in MD-901 cells (Figure S6B). Consistently, transcriptomic analysis of MD-901 CBC clones after three weeks of ibrutinib exposure revealed miR-28–dependent downregulation of both Akt2 and Erk2 (Figure S6B). Analysis of mTORC1 pathway activity by Western blot of characteristic phosphoproteins revealed that, in addition to pS6 (Ser240/244), miR-28 overexpression reduced phosphorylation of 4EBP1 (T37/46) in both MD-901 and Riva cells (Figure S6C). In Riva cells, we also observed a reduction of p70S6K phosphorylation at T389, a mTORC1-dependent modification that contributes to p70S6K activation and can promote S6 phosphorylation (Figure S6C). Consistent with these findings, miR-28 downregulated mTOR pathway genes in MD-901 cells that are also upregulated in ibrutinib-resistant ABC-DLBCL cell lines such as OCI-Ly1, OCI-Ly10, and HBL1(^31^) (Figure S6D). Overall, these findings show that miR-28 counteracts both mitochondrial and mTOR signaling adaptations associated with the development of ibrutinib resistance in ABC-DLBCL cells.

### Survival of ibrutinib-treated MCD DLBCL patients is associated with the miR-28–regulated ibrutinib resistance gene signature

To assess whether the transcriptional program induced by miR-28 in ibrutinib-treated ABC-DLBCL cells is associated with differences in survival among DLBCL patients, we analyzed outcome data from patients with the MCD (n=73), BN2 (n=31), and N1 (n=24) genetic subtypes of DLBCL who were treated with R-CHOP immunochemotherapy or R-CHOP plus ibrutinib and had five years of outcome data(^12^). Patients were ranked based on a singscore analysis(^32^) of the expression of genes associated with ibrutinib resistance from the miR-28-IR gene signature (logFC > 1), using baseline biopsies from the non-GCB DLBCL PHOENIX trial. Singscore is a method that assigns a score to each sample based on the relative expression of genes in a predefined signature, enabling comparison of signature activity across patients. Patients were then stratified into two groups (high and low score) according to the median score. Correlation with clinical outcomes revealed that MCD patients—the most prevalent genetic subtype related to ABC-DLBCL—with low singscore had higher overall survival than those with higher singscore when treated with ibrutinib, but not when treated with R-CHOP alone. However, this difference was statistically significant only in older MCD patients (age > 60 years; n = 54, p = 0.047) (Figure 4G, S6E). No significant differences in survival between patients with low and high singscore rankings were observed in N1 or BN2 DLBCL subtypes, regardless of ibrutinib treatment (data not shown).

Finally, in the cohort of older MCD patients (age > 60 years), we calculated a novel transcriptional score based on genes functionally identified as essential for the acquisition of ibrutinib resistance through a genome-wide CRISPR-Cas9 screen in two ABC-DLBCL cell lines (HBL1 and TMD8)(^20^). Using singscore, we assessed whether this ibrutinib-resistance score correlated with the miR-28–regulated IR gene signature identified in our study. Notably, we observed a positive and statistically significant correlation between the ibrutinib-resistance score and the miR-28–regulated IR signature score, driven by genes both regulated by miR-28 and identified as required for ibrutinib resistance acquisition in our study (Figure 4H). These results indicate that tumors with higher miR-28–associated transcriptional activity exhibit transcriptional features linked to a functional sensitivity to ibrutinib.

Collectively, these findings establish the miR-28 axis as a mechanistic regulator of transcriptional programs that drive ibrutinib resistance and influence clinical outcome in ibrutinib-treated MCD DLBCL patients.

### miR-28 suppresses the emergence and growth of ibrutinib-resistant ABC-DLBCL tumors

To evaluate the anti-tumor activity of miR-28 in ibrutinib-resistant B-cell lymphoma, we used different models of ABC-DLBCL with acquired resistance to ibrutinib. We first assessed the growth capacity of parental and ibrutinib-resistant U2932, MD-901, and Riva ABC-DLBCL cell lines, and found that the resistant cells exhibited enhanced growth in xenograft tumor models (Figure 5A). Next, we investigated whether miR-28 expression could inhibit the development of ibrutinib-resistant tumors in human ABC-DLBCL xenografts. Riva cells transduced with either scramble or miR-28-pTRIPZ constructs and Venus or Cerulean proteins were co-cultured *in vitro* for three weeks in the presence of doxycycline and progressively increasing concentrations of ibrutinib (ranging from 10-fold below to 2.5-fold below the GR_50_). After this period, RFP^+^ miR-28-Cerulean^+^ and RFP^+^ scr-Venus^+^ cells were sorted by flow cytometry and injected subcutaneously into NOD scid gamma (NSG) mice. The animals received doxycycline in their drinking water and daily doses of vehicle or 20 mg/kg ibrutinib, a dose that, as shown in our results, effectively inhibits parental Riva xenograft growth (Figure S7A). Scramble tumors in ibrutinib-treated mice were larger than those in vehicle-treated controls, indicating the emergence of *in vivo* ibrutinib resistance (Figure 5B). Notably, in ibrutinib-treated groups, xenografts with induced miR-28 expression exhibited significantly reduced tumor volumes compared to scramble controls (Figure 5B). These findings demonstrate that miR-28 inhibits the development of ibrutinib-resistant ABC-DLBCL tumors in xenografts.

**Figure 5.**
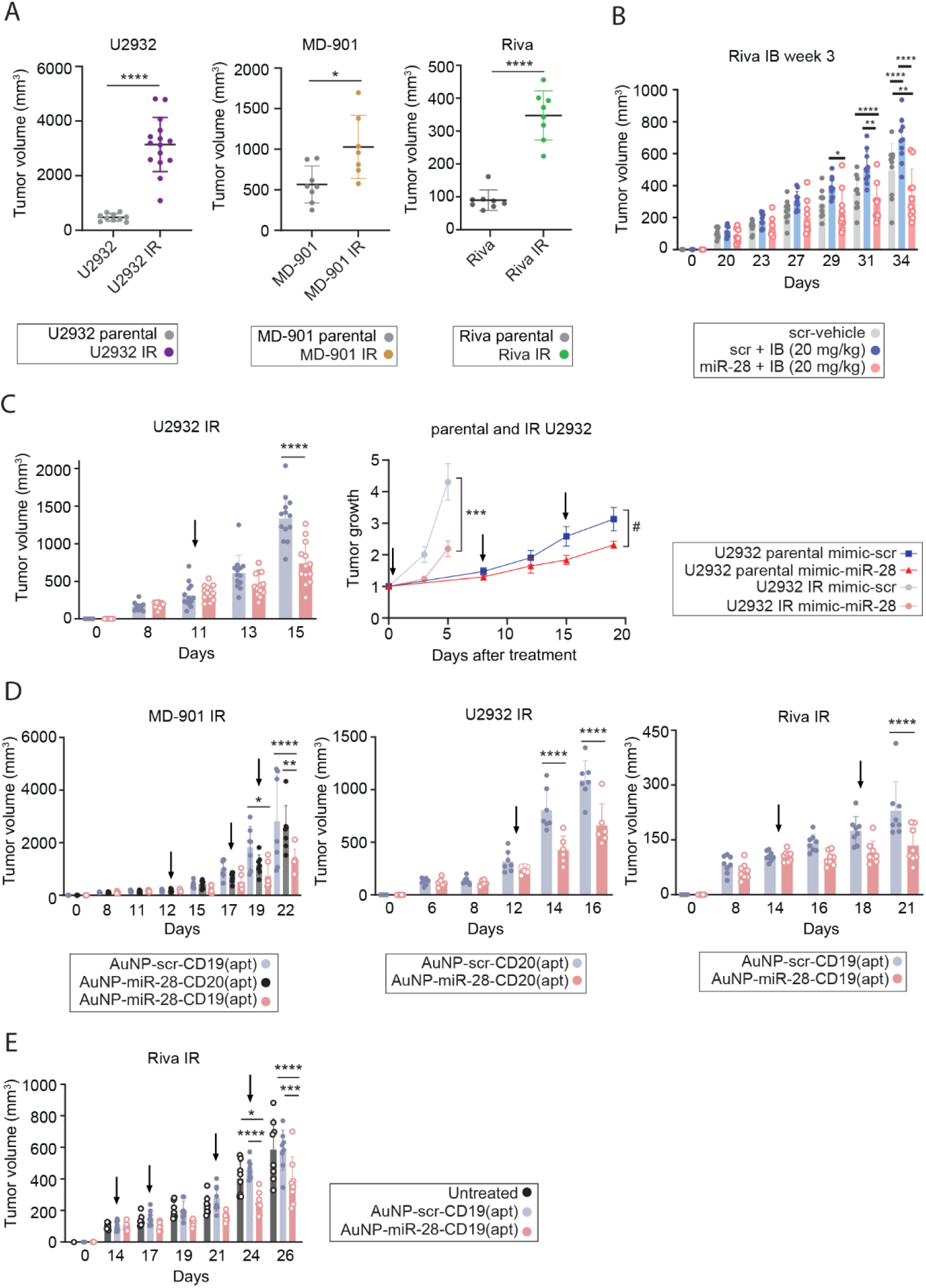
miR-28 suppresses the emergence and growth of ibrutinib-resistant DLBCL tumors in xenograft models. **(A)** Tumor volume of parental and ibrutinib-resistant ABC-DLBCL cell lines: U2932 (16 days post-injection; parental: n = 10, IR: n = 15), MD-901 (17 days post-injection; parental: n = 8, IR: n = 7), and Riva (23 days post-injection; parental: n = 8, IR: n = 8) in xenograft models using NOD scid gamma (NSG) mice. Data represent mean tumor volume ± SD. **(B)** Tumor volumes of scramble (scr; control) and miR-28–expressing Riva cell xenografts in NSG mice treated or untreated with ibrutinib. Scr-Venus and miR-28–Cerulean transduced Riva cells were co-cultured *in vitro* for 3 weeks in the presence of doxycycline and increasing concentrations of ibrutinib. RFP^+^ live cells were sorted and directly injected subcutaneously into NSG mice without further *in vitro* culture. The mice then received daily ibrutinib (20 mg/kg) or vehicle via oral gavage, and doxycycline in drinking water. Graph shows tumor volumes: scr tumors in vehicle-treated mice (gray dots, n = 10), scr tumors in ibrutinib-treated mice (blue dots, n = 10), and miR-28 tumors in ibrutinib-treated mice (pink dots, n = 10). Data represent mean ± SD. **(C)** Tumor volume of IR and parental U2932 xenografts in NSG mice treated intratumorally with miRNA mimics—miR-28 (light pink, n = 13) or scramble control (light blue, n = 13). Left graph: tumor volumes of U2932 IR xenografts. Data represent mean tumor volume ± SD. Right graph: tumor growth after treatment of U2932 IR (miR-28: light pink, n = 13; scr: light blue, n = 13) and parental U2932 xenografts (miR-28: red, n = 9; scr: dark blue, n = 10). Arrows indicate dosing schedules. Data represent mean ± SEM. **(D-E)** Tumor volumes of IR ABC-DLBCL xenografts treated intratumorally (D) or intravenously (E) with CD19- or CD20-targeted gold nanoparticles (AuNPs): MD-901 IR (scr-CD19: n = 7; miR-28-CD20: n = 7; miR-28-CD19: n = 6), Riva (intratumoral/intravenous: scr-CD19: n = 8/8; miR-28-CD19: n = 8/8), and U2932 (scr-CD20: n = 6; miR-28-CD20: n = 7). AuNP–miR-28 with cell line–specific aptamers (apt) (light pink), AuNP–miR-28 with non-specific apt (black), and AuNP–scramble with specific apt (light blue). Data represent mean ± SD. Arrows indicate dosing schedules. Statistical analysis: unpaired t-test (A); one-way ANOVA (B, C left graph, D and E); two-way ANOVA (C right graph). # or \**P* < 0.05; \*\**P* < 0.01; \*\*\**P* < 0.001; \*\*\*\**P* < 0.0001.

We further evaluated the therapeutic potential of miR-28 analogs delivered via different methods *in vivo* in models of ibrutinib-resistant ABC-DLBCL. First, ibrutinib-resistant U2932 ABC-DLBCL tumors were treated with synthetic miR-28 mimics or scramble controls. Once tumors reached at least 150 mm³, mice received intratumoral injections of 0.5 nmol of either miR-28 or control mimics. Treatment with miR-28 significantly inhibited the growth of ibrutinib-resistant U2932 tumors (Figure 5C). Notably, the inhibitory effect of miR-28 was stronger in ibrutinib-resistant tumors than in those derived from parental U2932 cells (2.1-fold reduction after one treatment vs. 1.3-fold reduction after three treatments, respectively; Figure 5C). Finally, we assessed whether aptamer-conjugated, lymphoma-targeted AuNPs carrying miR-28 could inhibit tumor growth. Ibrutinib-resistant ABC-DLBCL cell lines (U2932, MD-901, and Riva) were subcutaneously injected into NSG mice. After the tumors reached at least 150 mm³, mice were treated with intratumoral injections of control or miR-28 AuNPs conjugated to CD19- or CD20-specific aptamers. The xenografted cell lines expressed distinct surface markers (MD-901: CD19⁺CD20⁻; U2932: CD19^low^CD20⁺; Riva: CD19⁺CD20⁺; Figure S7B). Treatment of ibrutinib-resistant MD-901 tumors with CD19-targeted miR-28 AuNPs significantly inhibited tumor growth (Figure 5D, S7C-D). This antitumor effect was specific to miR-28 and the aptamer-targeted delivery, as the inhibition was observed compared to CD20-targeted miR-28 AuNPs or control AuNPs. Similar results were observed in ibrutinib-resistant U2932 and Riva xenografts treated with aptamer-conjugated miR-28 AuNPs targeting their respective surface markers (Figure 5D). We then assessed the efficacy of aptamer-guided miR-28 AuNPs following systemic administration. Ibrutinib-resistant Riva xenografts were established as in previous experiments, and mice were treated with twice-weekly intravenous injections of CD19-aptamer–conjugated miR-28 AuNPs. Mice receiving intravenous miR-28 AuNPs developed significantly smaller tumors than untreated mice or mice treated with control AuNPs (Figure 5E). Our findings demonstrate that ibrutinib-resistant tumors retain sensitivity to miR-28-mediated inhibition. Targeted delivery via aptamer-conjugated gold nanoparticles administrated locally or systemically enables this effect *in vivo*, underscoring the translational potential of miR-28 as a therapeutic strategy to counter resistance in ABC-DLBCL.

## DISCUSSION

In spite of the major advances in the identification of molecular drivers in DLBCL, a significant subset of these aggressive B-cell lymphomas remains a major clinical challenge(^3, 4^). Although ABC-DLBCL initially responds to BTK inhibitor–based targeted therapies, the emergence of acquired resistance continues to limit durable clinical benefit(^10, 14, 19^). In this study, we investigated whether the microRNA miR-28 could impair the development of non-genetic resistance to ibrutinib in ABC-DLBCL. We hypothesized that the activity of miR-28, which simultaneously inhibits the expression of several genes identified as essential for oncogenic signaling—including genes from the BCR-dependent PI3K and NF-κB pathways(^28^)— could disrupt the feedback activation and transcriptional rewiring associated with resistance.

Using a combination of flow cytometry–based competition assays, clonal barcoding with multicolor labeling, transcriptomic profiling, and *in vivo* xenograft models, we show that miR-28 impairs the emergence of ibrutinib-resistant cells in ABC-DLBCL. These results build upon seminal work identifying the transcriptional remodeling associated with the development of ibrutinib resistance in ABC-DLBCL, as well as the genes essential for the survival of resistant cells, as revealed by genome-wide CRISPR-Cas9 screens(^20, 21^). In ibrutinib-exposed MD-901 clones, miR-28 downregulated genes previously shown to be critical for resistance acquisition in HBL1 and TMD8 ABC-DLBCL cells(^20^). Pathway analysis of these clones further revealed that miR-28 suppresses the upregulation of mTOR and mitochondrial signaling—responses specifically induced after three weeks of adaptation to increasing ibrutinib doses. These observations are consistent with previous reports implicating PI3K/AKT/mTOR pathway activation in other ibrutinib- and PI3Kβ/δ inhibitor-resistant ABC-DLBCL models(^20, 31, 33^). Moreover, our reanalysis of publicly available datasets from resistant models(^31^) confirmed the enrichment of mitochondrial-related pathways, including those linked to resistance in HBL1, OCI-Ly10, and OCI-Ly1 cell lines. To support these findings, we assessed mTOR activity and mitochondrial function—such as ATP production rate and mitochondrial mass—in three ABC-DLBCL cell lines exposed to increasing ibrutinib concentrations, with or without miR-28 expression. These results confirmed the capacity of miR-28 to modulate key transcriptomic and metabolic changes associated with resistance development. Nevertheless, we found that miR-28 regulates a broad set of genes in MD-901 cells exposed to increasing doses of ibrutinib for three weeks, many of which are not directly related to the generation of ibrutinib resistance. Notably—and somewhat unexpectedly—although miR-28 inhibits the expression of cell-cycle–related genes in MD-901 clones under these conditions, consistent with our previous findings showing that the combination of miR-28 and ibrutinib induces a specific transcriptional cell-cycle arrest program in DLBCL(^29^), we did not observe differential regulation of proliferation-associated pathways among the subset of miR-28–regulated genes linked to ibrutinib resistance in MD-901 clones. These results suggest that the capacity of miR-28 to restrain the emergence of ibrutinib resistance is not primarily mediated by direct control of cell proliferation, but rather by post-transcriptional mechanisms that interfere with the adaptive rewiring of B cell lymphoma survival pathways, including mTOR and mitochondrial signaling, that promote tolerance to BTK inhibition.

Reinforcing the functional relevance of these transcriptomic alterations, we found that the expression of resistance-associated genes downregulated by miR-28—comprising the miR-28-IR signature—correlates with survival in ibrutinib-treated MCD-subtype DLBCL patients aged over 60 years in the PHOENIX clinical trial cohort. This clinical correlation is particularly relevant given that MCD and N1—two of the three genetic subtypes related to ABC-DLBCL—are the most responsive to ibrutinib due to their reliance on the BCR–NF-κB signaling pathway, with MCD being substantially more prevalent than N1(^12^). Additionally, most ABC-DLBCL cases occur in patients older than 60(^12, 34^). Altogether, these findings highlight the potential of miR-28 to target resistance-associated transcriptional programs of clinical relevance.

To investigate how these changes contribute to clonal selection and resistance emergence, we developed a clonal color barcoding flow cytometry–based assay that enabled us to track the fate of individual MD-901 clones during ibrutinib exposure. Approximately one-fourth of the clones became resistant after five weeks of dose escalation, driven by transcriptional adaptations that included the rewiring of metabolism-related pathways and activation of alternative survival signaling. While this model robustly captures the generation of ibrutinib resistance and associated epigenetic and transcriptomic adaptations, a fraction of non-adapting, ibrutinib-sensitive cells persist at the end of the 5-week ibrutinib exposure. Importantly, in cultures expressing miR-28, this clonal selection process was disrupted, with miR-28 acting as a brake on the emergence of resistant subpopulations. Nevertheless, a small fraction of RFP^+^ miR-28^+^ cells remained viable after the 5-week process in cell competition assays. Longer culture periods in the presence of 2x GR_50_ ibrutinib would be required to determine whether these cells are completely eliminated or have become tolerant to ibrutinib through activation of drug-tolerance mechanisms. Beyond *in vitro* analyses, we evaluated the development of ibrutinib resistance *in vivo* and demonstrated that miR-28 impairs the growth of ibrutinib-resistant xenografts.

While previous studies have proposed targeting individual proteins within alternative signaling pathways—such as components of the PI3K/AKT axis—to inhibit the growth of ibrutinib-resistant ABC-DLBCL cells or xenograft tumors(^31, 35–38^), whether resistance itself can be proactively prevented has remained unaddressed. Here, we demonstrate for the first time that the emergence of resistance in ABC-DLBCL can be actively suppressed—rather than treated only after it is established—using a miRNA-based intervention.

In addition to its preventive effects, we found that rapidly growing ibrutinib-resistant ABC-DLBCL cells remain sensitive to miR-28–mediated inhibition in xenografts. Notably, in our study, this effect was efficiently achieved through targeted delivery using aptamer-guided gold nanoparticles. By using aptamers tailored to each DLBCL cell line, AuNPs enabled efficient tumor targeting and uptake, as shown by the pronounced tumor suppression achieved with AuNP–miR-28–CD19 in MD-901 xenografts compared to non-specific controls. Such aptamer-guided AuNP delivery strategies, which can be administered locally or systemically, not only amplify on-target activity while minimizing exposure in non-malignant tissues, but also protect miR-28 from extracellular degradation and ensure sustained release. These properties highlight the clinical potential of aptamer-functionalized AuNPs as a selective and durable miRNA delivery system(^39–42^) for DLBCL treatment. Building on these findings, further studies focusing on: (i) the identification of DLBCL-specific cell surface molecules not expressed in non-malignant B cells and targetable with aptamers, (ii) the development of clinically scalable, tumor-specific miRNA-based drug delivery systems, (iii) the design of optimal miRNA-based combination therapies tailored to ABC-DLBCL patients across different genetic backgrounds and age groups, and (iv) the validation of these findings in clinical settings, will further enhance the translational potential of this work.

In summary, the results of this study provide insights into the development of targeted miR-28-based therapies delivered via aptamer-functionalized gold nanoparticles to prevent or overcome ibrutinib resistance in ABC-DLBCL.

## ACKNOWLEDGEMENTS

We thank Janssen and the European Genome-phenome Archive for providing access to data from the PHOENIX trial, the members of the B cell biology lab, Balbino Alarcón, José Luis Marina and María Jesús García de Yébenes for helpful discussions, and Arantzazu Alfranca for providing the LeGO vectors. E.A.C. and R.M.P were supported by PhD fellowships from the Comunidad de Madrid (PIPF-2022/SAL-GL-24469 and PIPF-2023/SAL-GL-29484 respectively), A.R.R. by CNIC funding and V.G.Y. by the Universidad Complutense de Madrid. B.S.E. is supported by a FPI contract from the Spanish Ministry of Science (PRE2022-102696). This work was funded by Spanish Ministerio de Ciencia e Innovación grants PID2019-107551RB-I00/AEI/10.13039/501100011033 and PID2022-137014OB-I00/AEI/10.13039/501100011033/ FEDER, UE to V.G.Y and PID2023-146982OB-I00 and CEX2020-001039-S to A.S and IMDEA Nanociencia, respectively. The CNIC is supported by the Instituto de Salud Carlos III (ISCIII), the Ministerio de Ciencia e Innovación (MCIN) and the Pro CNIC Foundation), and is a Severo Ochoa Center of Excellence (grant CEX2020-001041-S funded by MICIN/AEI/10.13039/501100011033). N.M.M. lab is supported by the grants CNS2022-135636, PID2021-126298OB-I00, PID2024-162320OB-I00 (founded by MCIU/AEI/10.13039/501100011033/FEDER, UE) and HR22-00447 (founded by La Caixa).

## AUTHOR CONTRIBUTIONS

E.A.C., R.M.P., M.L., U.M.P, A.D.B., T.F., B.S.E and S.M. performed experiments. C.G.E. performed computational analysis. M.M. synthesized aptamer- and miRNA-conjugated AuNPs. E.A.C., R.M.P., M.L., T.F., B.S.E and V.G.Y. analyzed and interpreted data. E.A.C. prepared figures and wrote Material and Methods. M.L, A.S., N.M.M. and A.R.R. contributed to the conception of the experimental design. V.G.Y. conceived the overall strategy and wrote the manuscript. All authors revised and contributed to the final manuscript.

---

**Figure S1.**
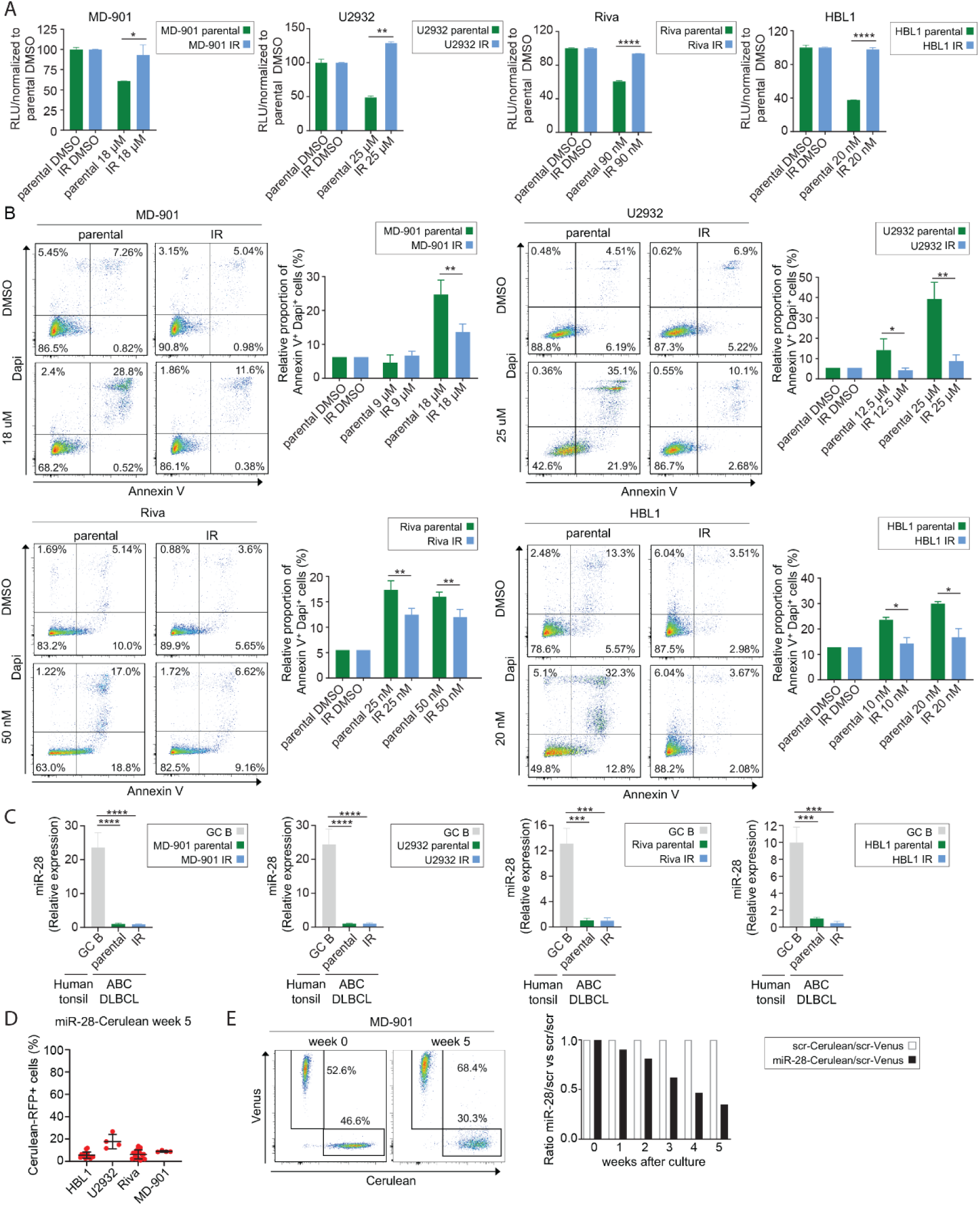
miR-28 suppresses ibrutinib resistance development in different ABC-DLBCL cell lines. **(A)** Relative number of live parental and ibrutinib-resistant (IR) MD-901, U2932, Riva, and HBL1 cells in DMSO- and ibrutinib-treated cultures. Cell viability was assessed three days after treatment with ibrutinib (IB) using the CellTiter-Glo® reagent, which quantifies total ATP. Data were normalized to DMSO-treated controls. **(B)** Cell death analysis by flow cytometry in parental and IR MD-901, U2932, Riva, and HBL1 ABC-DLBCL cell lines, analyzed after three days of culture with or without ibrutinib. Representative Annexin V and DAPI flow cytometry plots are shown (left). The proportion of dead cells (Annexin V⁺ DAPI⁺) in parental (green) and IR (blue) MD-901, U2932, Riva, and HBL1 cells is first normalized to the proportion of DMSO-treated cells and then expressed relative to the mean proportion of Annexin V⁺ DAPI⁺ cells in parental cells at the highest ibrutinib concentration for each line. **(C)** Quantification of miR-28 expression in parental and IR MD-901, U2932, Riva, and HBL1 ABC-DLBCL cell lines and in non-tumoral germinal center (GC) B cells isolated from human tonsils by qRT-PCR. Expression levels were normalized to the mean of the parental cell lines. **(D-E)** Polyclonal ibrutinib-resistance competition assays were performed by mixing ABC-DLBCL cells transduced with doxycycline-inducible RFP-expressing lentiviral vectors encoding either miR-28 or a scramble (scr) sequence and labeled with either Cerulean or Venus fluorescent proteins at a 1:1 ratio. Cultures were exposed to ibrutinib for five weeks, with the drug concentration increasing weekly to reach up to 2-fold above the ibrutinib GR_50_. (D) Quantification of miR-28-Cerulean RFP^+^ cells exposed to ibrutinib for five weeks in miR-28-Cerulean/scr-Venus competition assays of U2932 (n = 4), MD-901 (n = 4), Riva (n = 17) and HBL1 (n = 15) ABC-DLBCL cells. (E) Representative flow cytometry plots of scr-Cerulean/scr-Venus co-cultures and their evolution up to week 5 in MD-901 cells. The graph represents the growth ratio of miR-28-Cerulean/scr-Venus co-cultures and their evolution up to week 5, normalized to the week 0 ratio and then to scr-Venus/scr-Cerulean cells in the MD-901 DLBCL cell line. Values indicate mean ± SD (A-C). Statistical analysis: unpaired t-test (A-B); one-way ANOVA (C). *P < 0.05, **P < 0.01, *** P<0.001, ****P<0.0001.

**Figure S2.**
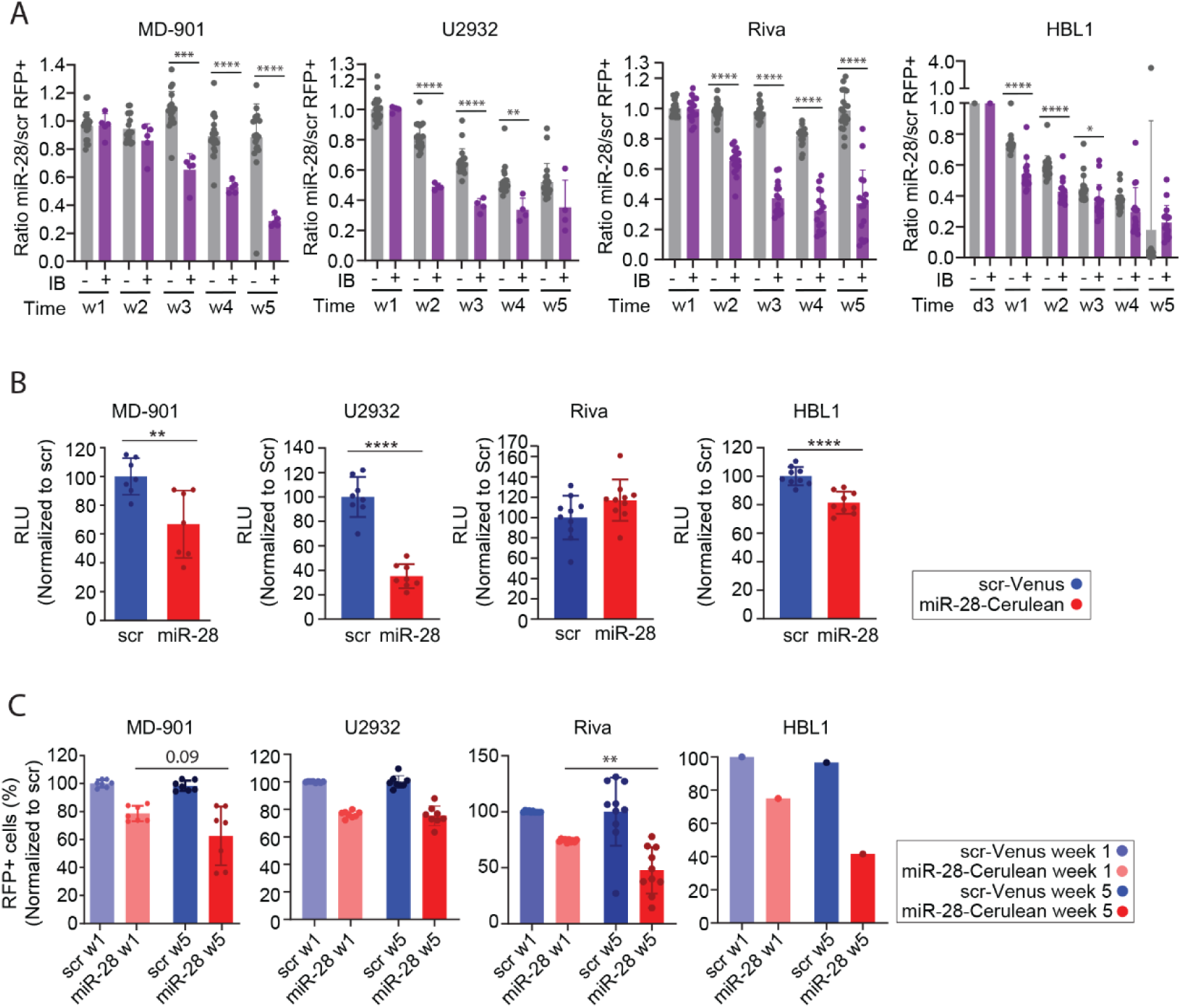
Enhanced miR-28–dependent decrease in cell fitness in cell competition assays in the presence of ibrutinib. **(A)** Polyclonal miR-28 competition assays were performed by mixing, at a 1:1 ratio, ABC-DLBCL cells transduced with doxycycline-inducible RFP-expressing lentiviral vectors encoding miR-28 and labeled with Cerulean, with cells transduced with scramble (scr) control vectors and labeled with Venus. Cultures were either left untreated or treated with ibrutinib for five weeks, with the drug concentration increasing weekly to reach up to 2-fold above the ibrutinib GR_50_. Graphs show the growth ratio of RFP⁺ miR-28-Cerulean to scr-Venus co-cultures and their evolution up to week 5, normalized to the week 1 ratio in MD-901, U2932, and Riva ABC-DLBCL cell lines, or to day 3 in HBL1 cells, under ibrutinib treatment (violet) or untreated conditions (grey). **(B-C)** Analysis of scr and miR-28 cultures grown for five weeks with increasing ibrutinib doses in four different ABC-DLBCL cell lines. (B) Viable cell number was determined at week 5 with CellTiter-Glo® reagent. (C) Graphs show the percentage of RFP^+^ cells at week 1 and week 5 in the presence of ibrutinib, analyzing the % RFP^+^ cells within each well normalized to the mean % RFP^+^ in scramble control wells at each corresponding time point: week 1 scr-RFP^+^ (light blue), week 1 miR-28-RFP^+^ (light red), week 5 scr-RFP^+^ (dark blue) and week 5 miR-28-RFP^+^ (dark red). Statistical analysis: upaired t-test (A and B); paired t-test (C). *P < 0.05, \*\**P*<0.01, *** *P*<0.001, \*\*\*\**P*<0.0001. Error bars denote SD.

**Figure S3.**
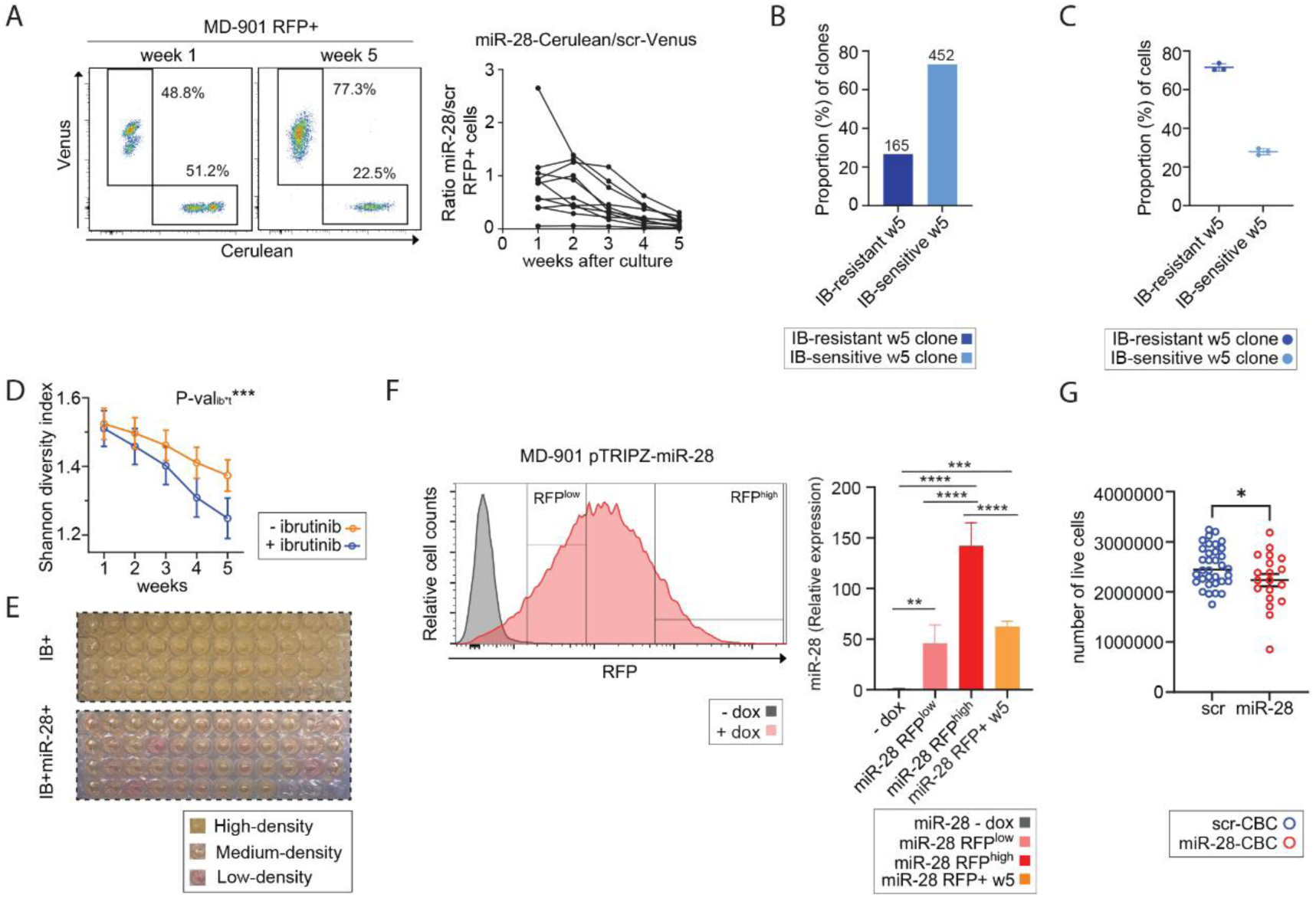
miR-28 impairs the development of ibrutinib resistance in ABC-DLBCL cells. MD-901 ABC-DLBCL cells were transduced with doxycycline-inducible, RFP-expressing lentiviral vectors encoding either miR-28 or a scramble (scr) sequence, and labeled with Cerulean and/or Venus fluorescent proteins. Cells were seeded at an average density of 10 cells per well and exposed to increasing concentrations of ibrutinib (IB), reaching up to 2-fold the GR_50_ after five weeks (A-E and G). Ibrutinib resistance miR-28-Cerulean versus scr-Venus competition assays performed are shown in (A), and color-barcoded MD-901 DLBCL (CBC) data are shown in (B-E and G). **(A)** Representative flow cytometry plots of RFP⁺ cells in scr–Venus/miR-28–Cerulean co-cultures in cell competition assays seeded at an average density of 10 cells per well over five weeks. The bar chart displays the growth ratio of miR-28–Cerulean to scr–Venus within the RFP⁺ population. **(B-C)** Proportion of ibrutinib-resistant (defined as week 5 > week 1) and ibrutinib-sensitive (defined as week 5 < week 1) clones (B) or cells (C) in scramble-transduced CBC culture wells after five weeks of culture, compared to week 1 (n = 3 experiments; 45 wells per experiment). **(D)** Clonal diversity measured by the absolute Shannon index in CBC culture wells treated with ibrutinib (blue) or left untreated (orange). **(E)** CBC wells after five weeks of ibrutinib exposure are classified as high-density, medium-density or low-density. Quantification is shown in Figure 3A. **(F)** Quantification of miR-28 expression in miR-28-pTRIPZ MD-901 ABC-DLBCL cells analyzed by qRT-PCR. RFP⁺ cells from miR-28-pTRIPZ MD-901 cultures were sorted according to their relative RFP expression levels after three days in the presence of doxycycline. Flow cytometry histograms showing RFP expression levels used to establish the sorting gates are shown. Cells without doxycycline (RFP^-^; -dox) were included as controls. miR-28 expression was also quantified in sorted RFP⁺ miR-28–Cerulean MD-901 cells from scr-Venus/miR-28–Cerulean cocultures after 5 weeks under escalating ibrutinib doses. Relative expression was normalized to the mean expression in -dox (RFP^-^) cells. Error bars denote SD of 3 independent wells. **(G)** Number of viable cells in scr (left) and miR-28–expressing (right) wells with more than 80% RFP⁺ cells after five weeks of culture, quantified using a Countess 3 Automated Cell Counter following trypan blue staining. Error bars represent SD in panels C-D and F, and SEM in panel G. Statistical analysis: two-way ANOVA (D), one-way ANOVA (F) and unpaired t-test (G). \**P* < 0.05, **P<0.01, *** P<0.001, ****P<0.0001.

**Figure S4.**
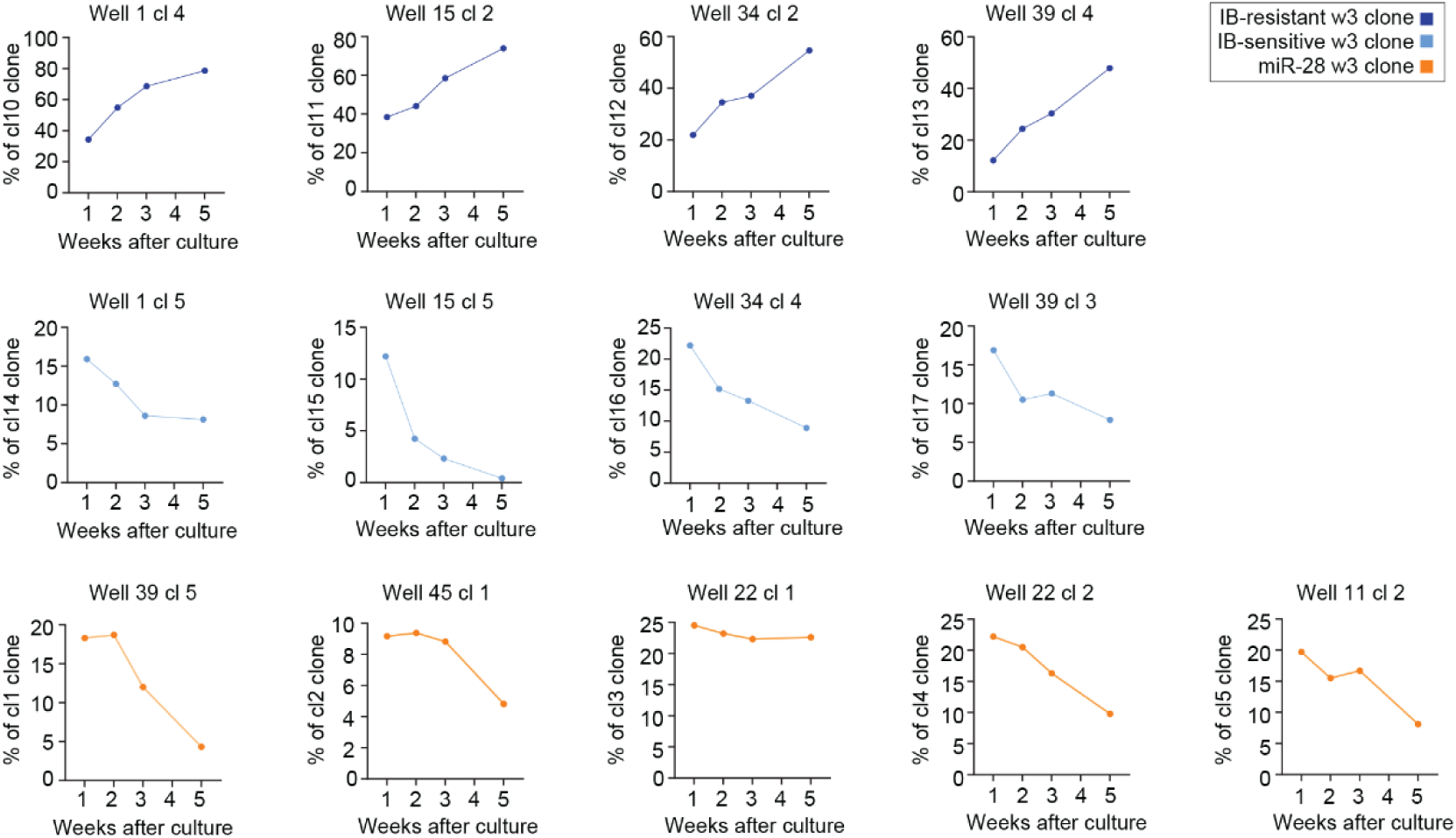
Evolution of the relative proportions of the scramble and miR-28 clones used in transcriptomic analysis. Graphs show the proportion of scr ibrutinib-sensitive clones, scr ibrutinib-resistant clones, and miR-28–expressing CBC clones used for RNA-seq after five weeks of ibrutinib exposure.

**Figure S5.**
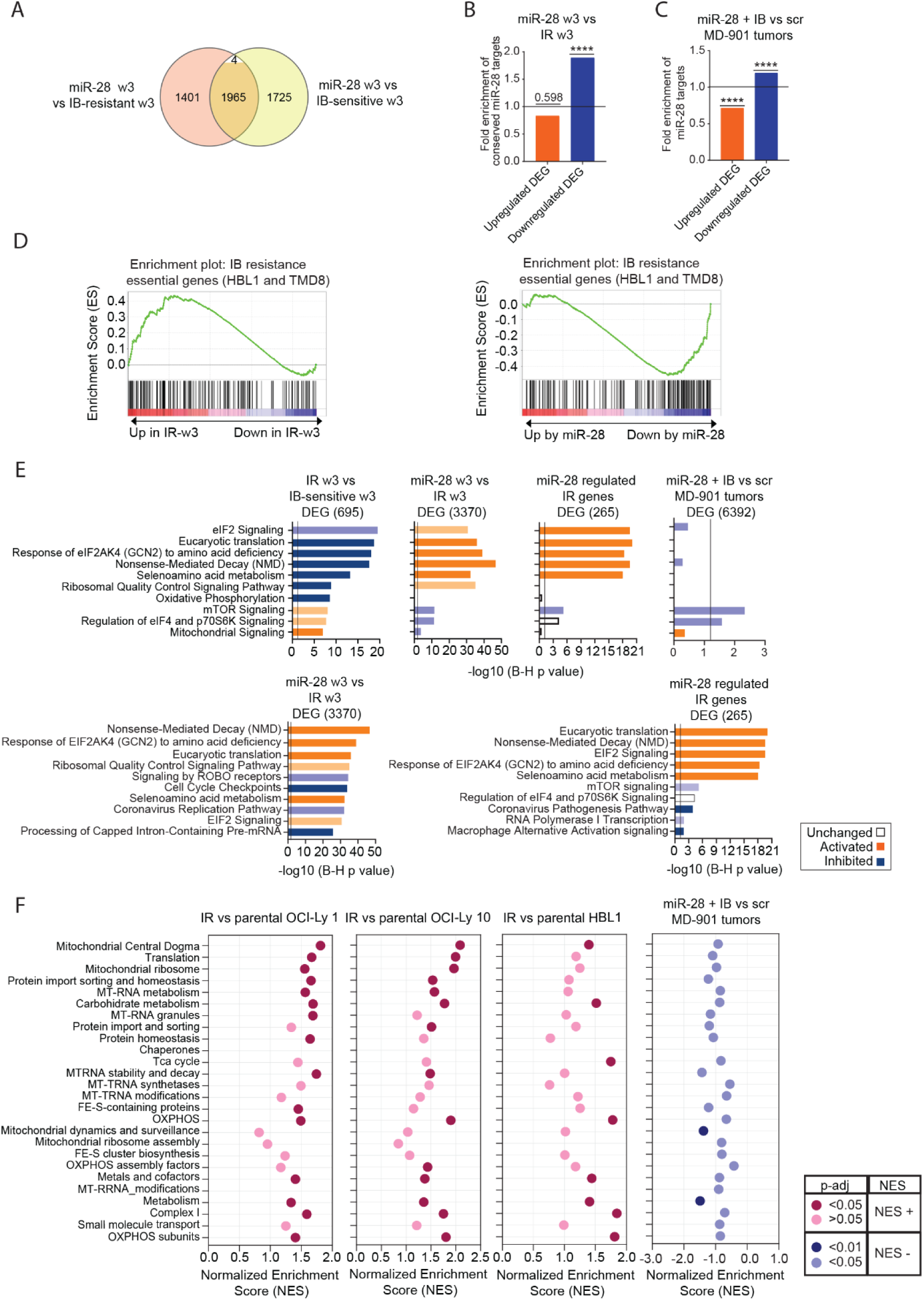
miR-28 regulates metabolic transcriptional programs associated with ibrutinib resistance. Transcriptomic analysis of sorted color-barcoded MD-901 (CBC) clones after three weeks (w3) of culture with gradually increasing concentrations of ibrutinib (IB) and of miR-28–expressing MD-901 tumors from ibrutinib-treated mice. **(A)** Venn diagram showing the number of DEGs between miR-28–expressing and scramble ibrutinib-resistant MD-901 clones (3370, shown in orange) and between miR-28–expressing and scramble ibrutinib-sensitive MD-901 clones (3694, shown in yellow) at week 3. DEGs shared between both comparisons (1969) are shown in the intersection and are colored when their expression changes are concordant (1965 genes, upregulated or downregulated by miR-28 in both comparisons) and shown in white when the direction of change differs (4 genes). **(B)** Fold enrichment of miR-28-predicted genes containing evolutionarily conserved miR-28 binding sites among transcripts differentially regulated by miR-28 in w3 CBC clones compared with IR w3 CBC clones, using all transcripts detected in the RNA-seq analysis as background. Two categories are shown: upregulated (odds ratio = 0.8; p = 0.5984) and downregulated (odds ratio = 2.2; p = 0.0002) transcripts. Statistical significance of enrichment was assessed using a χ² test with Yates’ correction. **(C)** Fold enrichment of miR-28–predicted targets among transcripts differentially regulated by miR-28 in MD-901 tumors from ibrutinib-treated mice compared with control MD-901 tumors from untreated mice, using all transcripts detected in the RNA-seq analysis as background(^9^).Two categories are shown: upregulated (odds ratio = 0.63; p = 0.0001) and downregulated (odds ratio = 1.33; p = 0.0001) transcripts. Statistical significance of enrichment was assessed using a χ² test with Yates’ correction. **(D)** Gene Set Enrichment Analysis (GSEA) of a BTK inhibitor resistance-essential gene set(^3^) in IR and miR-28 w3 CBC clones (FWER p = 0.009, FDR q = 0.009, ES = 0.43 in IR w3 vs. ibrutinib-sensitive w3; FWER p = 0.000, FDR q = 0.000, ES = -0.46 in miR-28 w3 vs. IR w3). **(E)** Graphs display the top 10 significantly altered pathways identified by Ingenuity Pathway Analysis (IPA). The comparison between ibrutinib-resistant and ibrutinib-sensitive w3 clones is shown in the top left panel. Pathway activity for the same pathways is shown for the comparison of miR-28 vs. ibrutinib-resistant w3 clones (top second from left), the miR-28–regulated IR gene signature (top third from left), and miR-28–expressing MD-901 tumors in ibrutinib-treated mice (top right).The bottom panels display the top 10 significantly altered pathways for the comparison between miR-28 vs. ibrutinib-resistant w3 clones (bottom left) and the miR-28–regulated IR gene signature (bottom right). Predicted pathway activity is indicated by color (blue: inhibited; orange: activated; white: non-predicted). P-values were adjusted for multiple testing using the Benjamini–Hochberg false discovery rate (FDR) method. **(F)** GSEA of mitochondrial-related pathways (Mitocarta 3.0) in ibrutinib-resistant (IR) versus parental OCI-Ly1, OCI-Ly10, and HBL1 cells(^6^) and in miR-28–expressing MD-901 tumors in ibrutinib-treated mice(^9^). Normalized enrichment scores (NES) and adjusted p-values (Benjamini–Hochberg correction) for significantly altered pathways are displayed in a scatter plot. Activated pathways are highlighted in pink and inhibited pathways are highlighted in blue.

**Figure S6.**
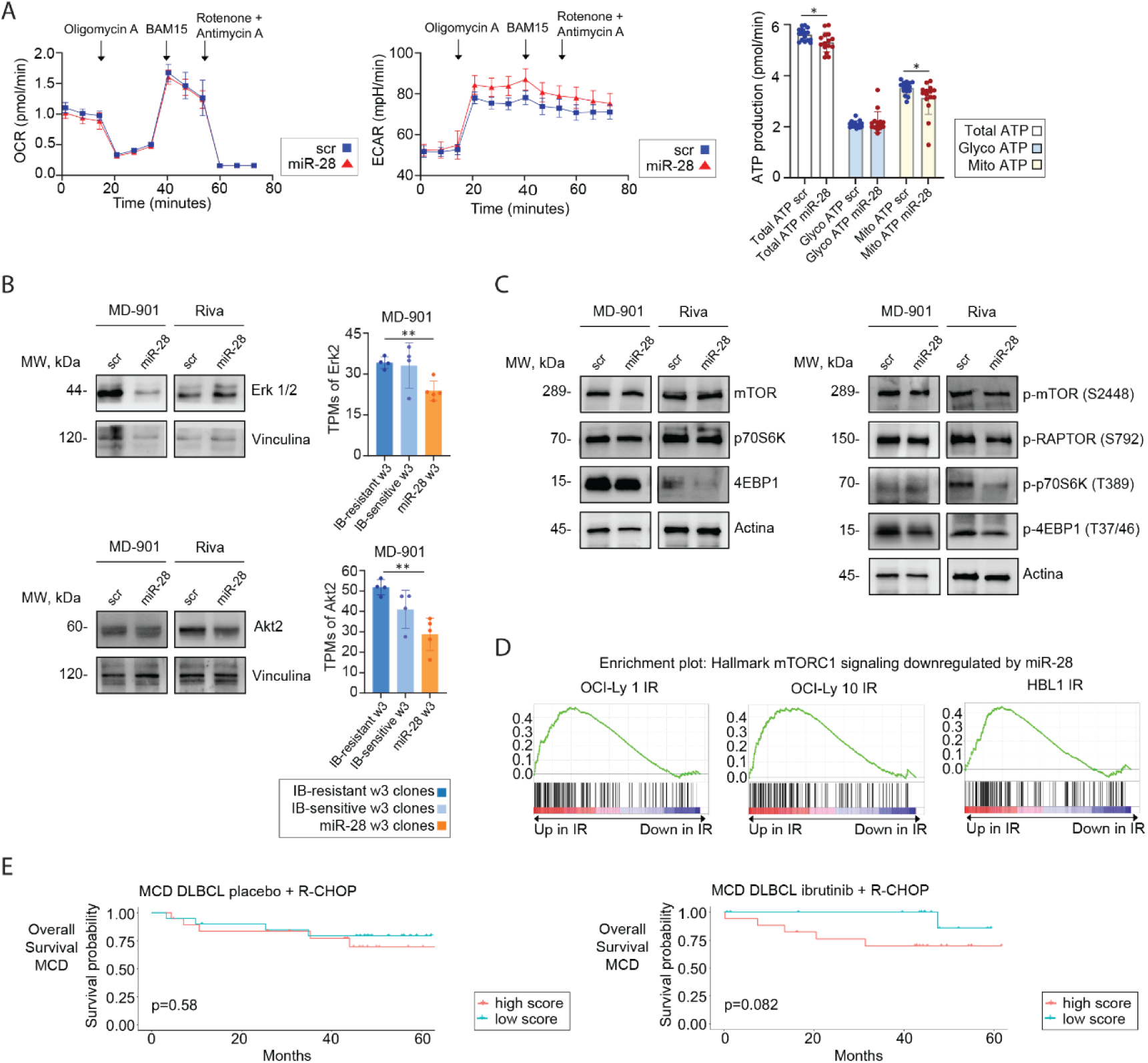
miR-28 regulates mTOR signaling and cellular ATP production. **(A)** Oxygen consumption rate (OCR) and extracellular acidification rate (ECAR) measurements in miR-28– and scr-expressing MD-901 cells after three weeks of culture in the presence of gradually increasing concentrations of ibrutinib (miR-28: red; scr: blue). OCR and ECAR were measured before and after sequential injections of oligomycin A, BAM15, and rotenone plus antimycin A. Arrows indicate the timing of each injection. The chart shows ATP production rates: total (white), glycolytic (blue), and mitochondrial (yellow). **(B-C)** Western blot and transcriptomic analyses of Erk2 and Akt2, predicted miR-28 targets upstream of mTOR signaling. Panel B includes Western blot and transcriptomic data, whereas panel C shows Western blot analysis of mTOR pathway activity based on phosphorylation of the indicated components. Western blots were performed on lysates from sorted RFP⁺ scramble and miR-28 pTRIPZ–transduced MD-901 and Riva ABC-DLBCL cells collected 48 h after doxycycline treatment. Transcriptomic data (panel B) show TPM of Akt2 and Erk2 in scramble ibrutinib-resistant (IR) w3 clones (dark blue, n = 4), scramble ibrutinib-sensitive w3 clones (light blue, n = 4), and miR-28–expressing CBC clones (orange, n = 5). **(D)** Gene Set Enrichment Analysis (GSEA) of the “Hallmark mTORC1 signaling” pathway (as defined by GSEA), showing downregulation by miR-28 in CBC MD-901 clones. Comparisons were made between IR and parental OCI-Ly1, OCI-Ly10, and HBL1 cells(^6^). Significant enrichment was observed in IR cells. **(E)** Kaplan–Meier analysis of overall survival in MCD DLBCL patients (n = 73), stratified by miR-28–IR gene signature score in baseline biopsies from the PHOENIX trial(^7^). Patients were treated with R-CHOP or R-CHOP plus ibrutinib. Groups correspond to the highest 50^th^ percentile (up, red) and lowest 50^th^ percentile (down, green). Statistical analysis: unpaired t-test (A and B); Log-rank (Mantel–Cox; E). \**P* < 0.05; \*\**P* < 0.01.

**Figure S7.**
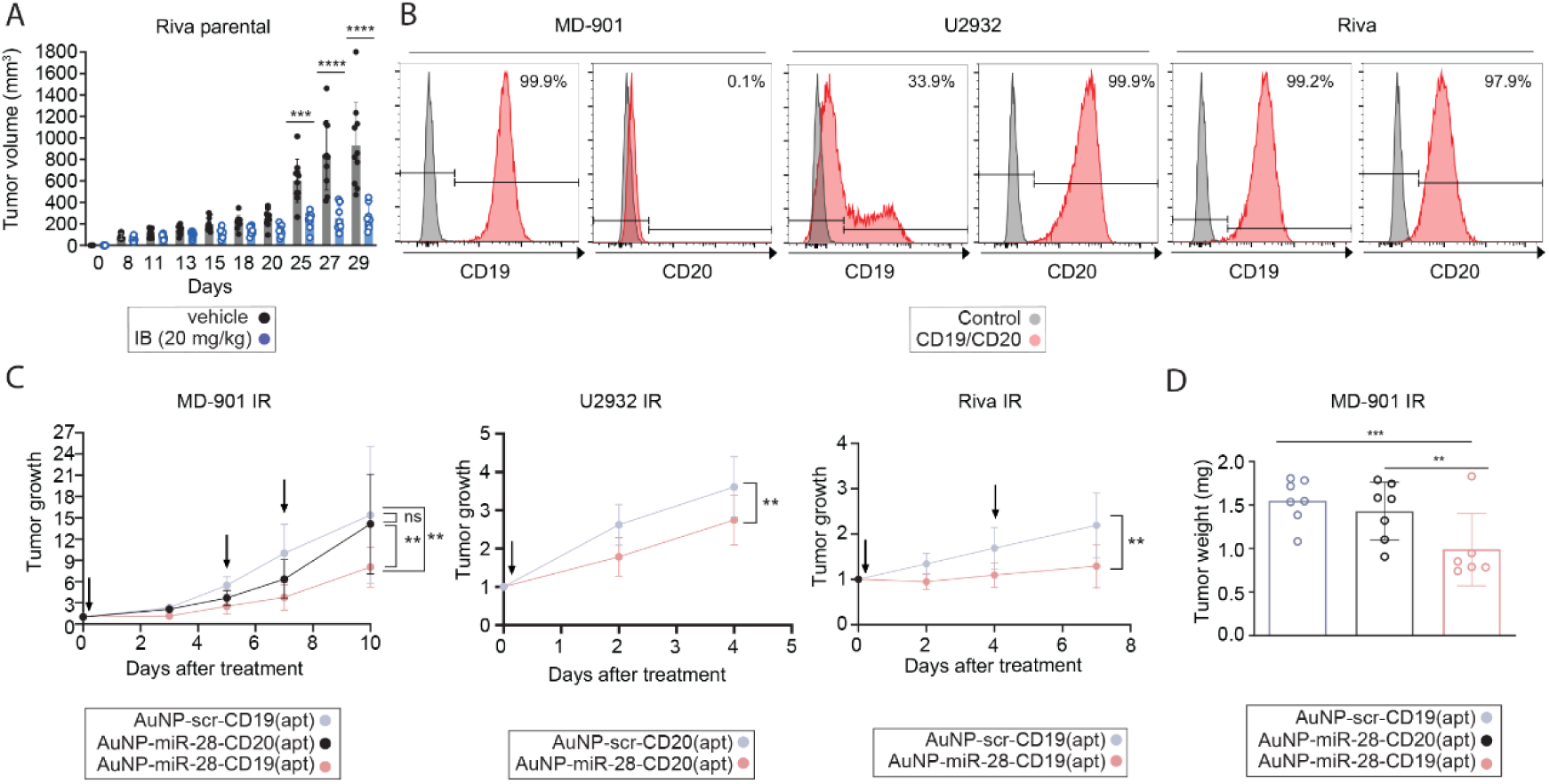
Aptamer-targeted delivery of miR-28 suppresses the growth of ibrutinib-resistant DLBCL tumors in xenograft models. **(A)** Tumor volume of parental Riva DLBCL cells in xenograft models in NSG mice treated daily with ibrutinib (20 mg/kg, n = 8) or vehicle (n = 10) via oral gavage. **(B)** CD19 and CD20 expression in MD-901, U2932, and Riva cell lines analyzed by flow cytometry. **(C–D)** Ibrutinib-resistant DLBCL cells were generated following the protocol described in Figure 1. (C) Tumor growth of ibrutinib-resistant (IR) ABC-DLBCL xenografts treated intratumorally with CD19- or CD20-targeted gold nanoparticles (AuNPs): AuNP–miR-28 with specific aptamers (light pink), AuNP–miR-28 with nonspecific aptamers (black), and AuNP–scr with specific aptamers (light blue). Arrows indicate dosing schedules. (D) Tumor weight measured at endpoint in IR MD-901 xenografts treated with: AuNP–scr–CD19 (light blue), AuNP–miR-28–CD20 (black), and AuNP–miR-28–CD19 (light pink). Statistical analysis: one-way ANOVA (A); two-way ANOVA (C); unpaired t-test (D). \*\**P* < 0.01; \*\*\**P* < 0.001; \*\*\*\**P* < 0.0001. Data represent mean ± SD.

